# Integrated Single-cell Analysis Uncovers Regulatory Logic of Cranial Ectoderm Development

**DOI:** 10.64898/2025.12.17.694990

**Authors:** Ceren Pajanoja, Jenaid Rees, Ed Zandro M Taroc, Laura Kerosuo

**Affiliations:** National Institute of Dental and Craniofacial Research, Intramural Research Program, Neural Crest Development and Disease Unit, National Institutes of Health, Bethesda, USA; University of Helsinki, Faculty of Medicine, Biochemistry and Developmental Biology Department, Helsinki, Finland

## Abstract

Embryonic ectoderm patterning in the cranial region establishes the neural plate, neural crest, sensory placodes, and non-neural ectoderm in coordination with the underlying mesoderm and endoderm. Although many studies have defined the signaling pathways and transcriptional regulators driving individual lineage specification, the temporal and spatial coordination among adjacent tissues remains poorly understood. To address this, we analyzed a single-cell RNA sequencing (scRNAseq) data set of a developmental series from the chick midbrain axial level, spanning stages from gastrulation to post-neurulation. Focusing on a defined region enabled high-resolution analysis of transcriptional dynamics and subpopulation transitions. Transcription factor downstream activity inference identified several ubiquitously expressed chromatin and histone modifiers—previously not associated with ectodermal patterning—that exhibit canonical downstream activity spatiotemporally restricted to specific developing domains. Moreover, we uncovered genes that remain canonical downstream activity throughout the progression of individual ectodermal domains, while others displayed negative activity scores, reflecting a repressed or alternatively a non-canonical downstream activity status that changed between cell types and developmental stage but was independent of transcription factor binding availability status on open chromatin. Finally, ligand–receptor interaction analyses across germ layers highlighted global signaling networks coordinating early embryogenesis, which were verified by *in situ* hybridization-based gene expression data. In sum, our study provides novel insights into the gene regulatory inputs that shape ectodermal cell fate decisions during neurulation and establishes a new framework for analyzing scRNAseq data to understand tissue patterning over developmental time.

## INTRODUCTION

After gastrulation, the ectoderm in the cranial region divides medially into the neural plate, which forms the central nervous system (CNS), and is neighbored by the neural plate border region that gives rise to the neural crest and adjacent sensory placode domains. The most lateral ectodermal territory becomes the non-neural ectoderm, which ultimately forms the skin (1–4). Similarly, the underlying mesoderm and endoderm layers are patterned into spatially and functionally distinct domains (5–8).

Extensive research over several decades has focused on identifying the inductive signals and downstream gene regulatory networks that drive fate specification and maturation of these domains (5, 8–21). Recent advances in RNA sequencing, multiplexed spatial transcriptomics, and imaging-based studies suggest that ectodermal patterning occurs gradually rather than through the sharp establishment of territories immediately after gastrulation (22–24). However, most existing data focus on a single domain (24, 25), neglecting the timing and interrelationships between neighboring tissues, which limits understanding of development *in vivo*.

To gain a more integrated view of embryogenesis, we collected a developmental series from a single axial level—the chick midbrain—and performed single-cell RNA sequencing (scRNAseq) to explore transcriptional changes (NCBI GEO database under accession number GSE221188). This approach addresses challenges in whole-embryo analyses, where subtle subpopulation differences are often obscured by presence of multiple axial levels, limitations in sample size and sequencing depth (26, 27). By focusing on a defined embryonic region, we achieved a higher-resolution view of transcriptional dynamics and subpopulation evolution over developmental time.

The advent of scRNAseq has transformed our understanding of transcriptional dynamics during development and tissue homeostasis. While RNA expression is often interpreted as a proxy for protein activity, various regulatory mechanisms can disrupt this correlation. RNA molecules can be inhibited (e.g., by siRNA, lncRNA, or RNA stability regulation), and protein function is further modulated by post-translational modifications. In the case of transcription factors (TFs), modifications in histone and DNA methylation and acetylation status alter chromatin accessibility and determine whether enhancer regions are available for binding. Consequently, high RNA expression does not necessarily indicate active protein function.

Single-cell multiomics approaches now allow comparisons between RNA expression and chromatin accessibility (e.g., ATAC-seq) to predict potential TF binding, and ChIP-seq to confirm enhancer occupancy. However, these analyses do not distinguish whether a bound TF acts as an activator or repressor or reflects an intermediate, poised status, leaving uncertainty about its downstream effects. Moreover, even multiomics studies remain incomplete, as they rarely capture the full range of post-translational modifications that affect protein activity. Due to resource constraints, many studies still rely on only one type of assay rather than integrating multiple modalities.

In this study, we analyzed a developmental series of cranial chick embryos at the midbrain level, spanning stages from immediately after gastrulation to post-neurulation. To achieve deeper insights into transcriptional regulation and gene regulatory networks, we applied **TF Activity Inference** (28) —a recently developed algorithm that distinguishes cells based on the canonical downstream activity of expressed genes. TF activity was calculated using the *decoupleR* framework and the **CollecTRI** resource (29) a comprehensive database of TF–target interactions compiled from twelve metadata sources, including public databases (e.g., DoRothEA, SIGNOR), text mining (ExTRI), manual curation, and regulatory annotations. Rather than relying solely on RNA levels, this approach infers TF activity based on the expression of their canonical target genes, providing a more functional perspective on regulatory dynamics derived from scRNAseq data.

Additionally, we investigated regulatory signaling interactions between developing cell types, including those in the underlying mesoderm and endoderm. Embryogenesis is orchestrated by the interplay of activating and inhibitory signals across a few key signaling pathways (30–33). Numerous studies have demonstrated that Bmp activation versus inhibition distinguishes non-neural ectoderm from neural plate (31, 34–36), that Fgf signaling inhibits Bmp to induce neural fate (13, 37) and that Wnt signaling promotes neural plate border formation (38–40). Despite this, the precise sources and dynamics of signaling molecules remain unclear, as most studies rely on ectopic manipulations. To address this, we performed **CellChat** analysis (41, 42) to identify ligand–source relationships underlying key signaling events, which provided information novel interactions and source tissues.

In summary, our study provides an integrated framework for analyzing developmental dynamics at single-cell resolution, offering new insights into transcriptional regulation, signaling interactions, and analytical strategies to extract greater value from existing scRNAseq datasets.

## Results

### Ectodermal subsets represent different cell type domains

To explore transcriptional changes during early head development, we investigated time points ranging from immediately post gastrulation to post neurulation stage. ScRNAseq analysis was performed for a total of 28,267 cells from the chicken embryo midbrain across five developmental time points – HH5, 1 somite, 4 somites, 7 somites (22), and 13–14 somites (HH11) (**Fig. 1A**). Each sample was individually preprocessed, and validations confirmed the absence of batch effects (**Supplemental Fig. 1A-C**). For each individual embryonic stage, cells clustered according to germ layers; the endoderm, mesoderm (with a separate notochord), and ectoderm (**Supplemental Table 1**). For deeper understanding of ectodermal transcriptional dynamics, we further subsetted the ectoderm, which divided into clusters of non-neural ectoderm, neural plate border, neural crest, ventral neural tube and rest of the neural tube, which included the dorsal and lateral parts. All subpopulations expressed typical genes of the respective cell type, as highlighted in **Fig. 1A**. Additionally, in line with our previous findings (22), during the three youngest stages, we identified a large population of “undecided” pan-ectodermal stem cells that expressed stem-cell related genes like Sall4, Sall1, Pou5f3, Lmo1, Cdk1, Hesx1 and Tubb3 (43, 44) and lower levels of both neural and non-neural markers (Nestin, Hes4, Dlx5, Cldn1, Krt24). Furthermore, cells of the neural plate border (where neural crest is induced) emerged at mid-neurula stage, and neural crest populations were detected at the latest time points (**Fig. 1A, and Supplemental Table 1**).

**Figure 1.**
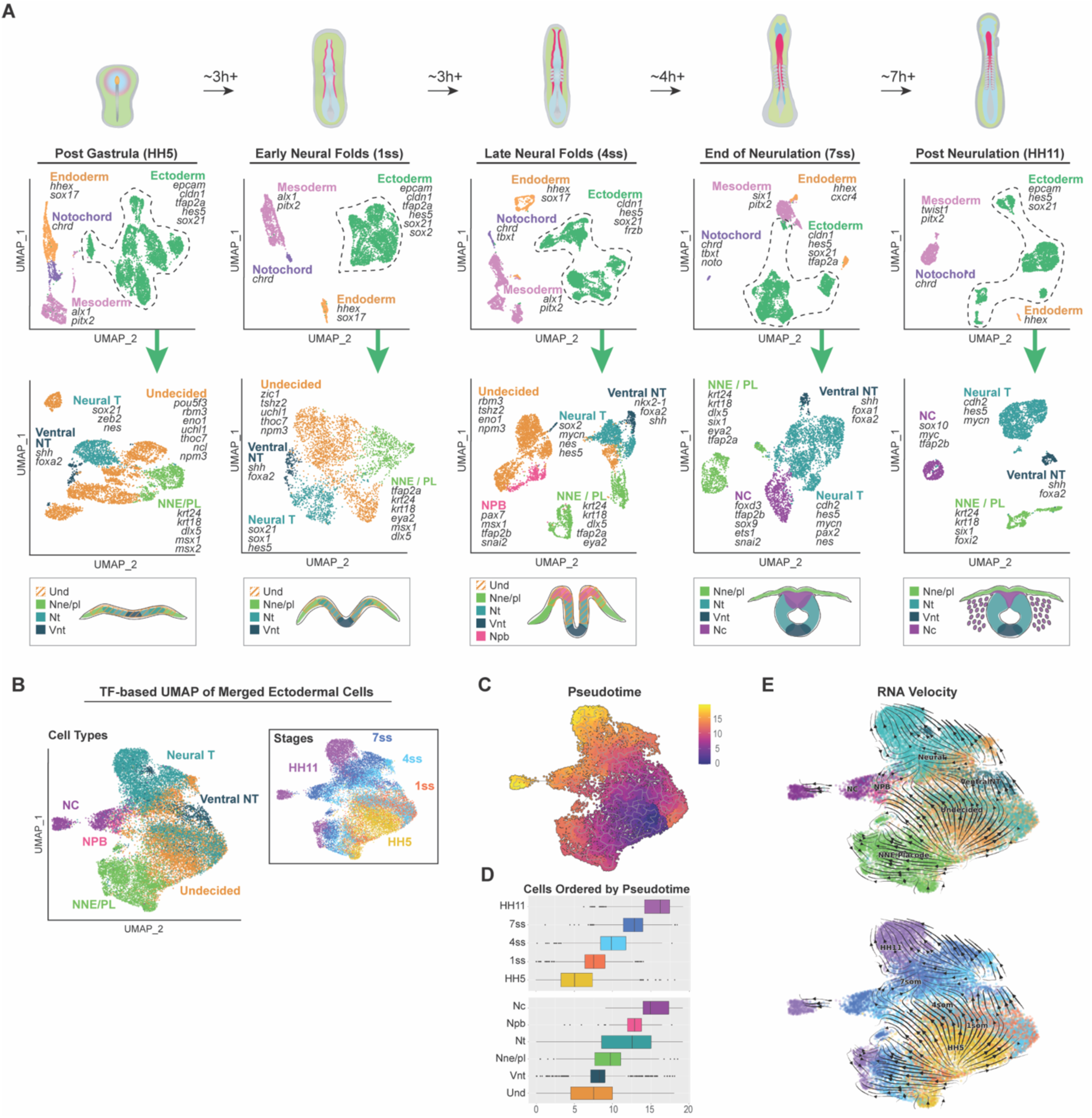
Ectodermal subpopulations stem from undecided pan-ectodermal stem cells. **A**) scRNAseq dataset was collected from chick embryos at midbrain level, spanning stages from right after gastrulation (HH5) to post-neurulation (HH11). Stage-specific UMAPs show segregated cells into three germ layers based on transcriptional profiles (upper panel). The ectoderm was further subsetted into subdomains, and key differentially expressed genes are highlighted for each group. **B)** UMAP, constructed by only using transcription factor expression, illustrates the distribution of ectodermal cell types and their developmental stages. **C)** Pseudotime trajectory overlaid on the transcription factor-based UMAP shown in panel B (Trajectory inference was performed using the full gene expression matrix). **D)** Distribution of pseudotime values across developmental stages and ectodermal cell types, shown as box plots. **E)** RNA velocity analysis shows that all ectodermal cell types stem from the undecided pan-ectodermal stem cells (direction of arrows).

### Ectodermal subdomains stem from undecided pan-ectodermal stem cells in a gradual process

Traditionally the ectoderm is thought to be patterned into the non-neural, future epidermis, neural crest and neural domain of the central nervous system immediately after gastrulation (45). Our recent finding of the entire ectoderm maintaining a large subset of cells with a pluripotency-like transcriptional profile doesn’t support this hypothesis but suggests that decision making behind ectodermal fates is a much more gradual process (22), which also is in line with two other recently published scRNAseq analyses (24). On the other hand, different hypotheses have also been proposed: a fairly recent study suggests that neural crest cells are reprogrammed to a pluripotent-like state in the ectoderm at mid-neurula stage after the ectoderm having first lost its pluripotency during gastrulation (46), and one of the earliest whole embryo scRNAseq studies in zebrafish embryo also came to a conclusion that neural crest cells may not be arising from the neural plate border region (47), thus building controversy on the topic on whether or not the entire ectoderm stems from a single, early pan-ectodermal population. We asked the question if the undecided, pan-ectodermal stem cells identified in our axial level specific data set, which share a transcriptional profile with all ectodermal domains by expressing lower levels of genes specific to each of the respective domain, have a transcriptional profile that places them as the initial stem cell population for all ectodermal subpopulations. For this, to enable trajectory inference across the entire ectodermal dataset, we aimed to construct a continuous UMAP embedding that would represent developmental transitions from early to late stages. However, when using the full gene set, the initial UMAP layout resulted in disjoint clusters with limited connectivity across developmental stages, preventing reliable pseudotime or RNA velocity analysis (**Supplemental Fig. 1D**). Given that transcription factors (TFs) are central regulators of cell fate decisions and are more likely to capture underlying regulatory dynamics, we redefined the dimensionality reduction using only TFs. We generated a curated list of TF genes based on chick and human annotations (sourced from KEGG BRITE), and subsetted our data accordingly. Recomputing the UMAP using this TF-only expression matrix produced an improved embedding, with cell populations from all five developmental stages aligning along a continuous manifold, enabling stage-spanning visualization of developmental relationships (**Fig. 1B).** To reconstruct developmental trajectories, we applied pseudotime analysis using Monocle3 (48), initializing the trajectory from the earliest stage (HH5) (**Fig. 1C**). While the UMAP was constructed using TF expression only, we reintroduced the full gene expression matrix for trajectory inference and subsequent analyses. The pseudotime ordering revealed a smooth progression aligned with developmental stage, with early cells positioned at one end and late-stage populations at the other (**Fig. 1D**). We then examined pseudotime distributions across stages and cell types using box plots, which showed that ventral neural tube cells traverse the shortest developmental path, while neural crest cells exhibit the longest pseudotime span, consistent with a more extended and plastic developmental trajectory (**Fig. 1D**). To further assess developmental directionality, we applied RNA velocity analysis using the full transcriptome and projected the resulting velocity vectors onto the TF-based UMAP (49). The velocity fields, based on proportions of spliced vs unspliced transcripts of all respective genes within individual cells (50), revealed a continuous temporal progression from HH5 to HH11, with trajectories demonstrating that ectodermal fates, including the neural crest cells that transit through the neural plate border stage, emerge from an initially uncommitted pan-ectodermal stem cell population (**Fig. 1E**), in agreement with the pseudotime results and previous findings (22).

### TF Activity Inference Reveals Dynamic Regulation Across Stages and Cell Types

We next focused on identifying transcriptional regulators underlying ectodermal patterning. To this end, we performed differential expression (DE) analysis restricted to TF genes. DE analysis was conducted between cell types, and the top differentially expressed TFs from the analysis were visualized in a heatmap (**Fig. 2A, Supplemental Table 2**), revealing both temporal and spatial regulators that included several known transcription factors previously described in ectodermal gene regulatory networks. Next, we calculated the canonical TF downstream activity using the decoupleR framework and the CollecTRI resource, a curated database of TF–target interactions (**Fig. 2B**) (28). Rather than relying solely on RNA expression, this method calculates TF activity based on the expression of their predicted target genes, providing a more functional perspective on regulatory dynamics. Because TF activity inference requires information from downstream genes rather than the TFs themselves, we generated a higher-resolution set of biological groups by combining developmental stage and cell type into 21 distinct stage-specific cell type identities. This approach ensured that the input to the activity analysis reflects stage-specific regulatory changes rather than signals averaged across stages or cell types. We then applied decoupleR to log fold change (LFC) values obtained from differentially expressed genes across these 21 unique identities (**Supplemental Fig.2A, Supplemental Table 2**), as LFC emphasizes biologically meaningful changes in expression while minimizing the influence of baseline expression levels. Next, to explore the regulatory relevance of the same TFs that were highlighted based on RNA expression level differences (**Fig. 2A**) we examined their inferred activity scores and, interestingly, noticed that canonical downstream activity varied by gene irrespective of RNA expression level (**Fig. 2A compared to 2C**).

**Figure 2.**
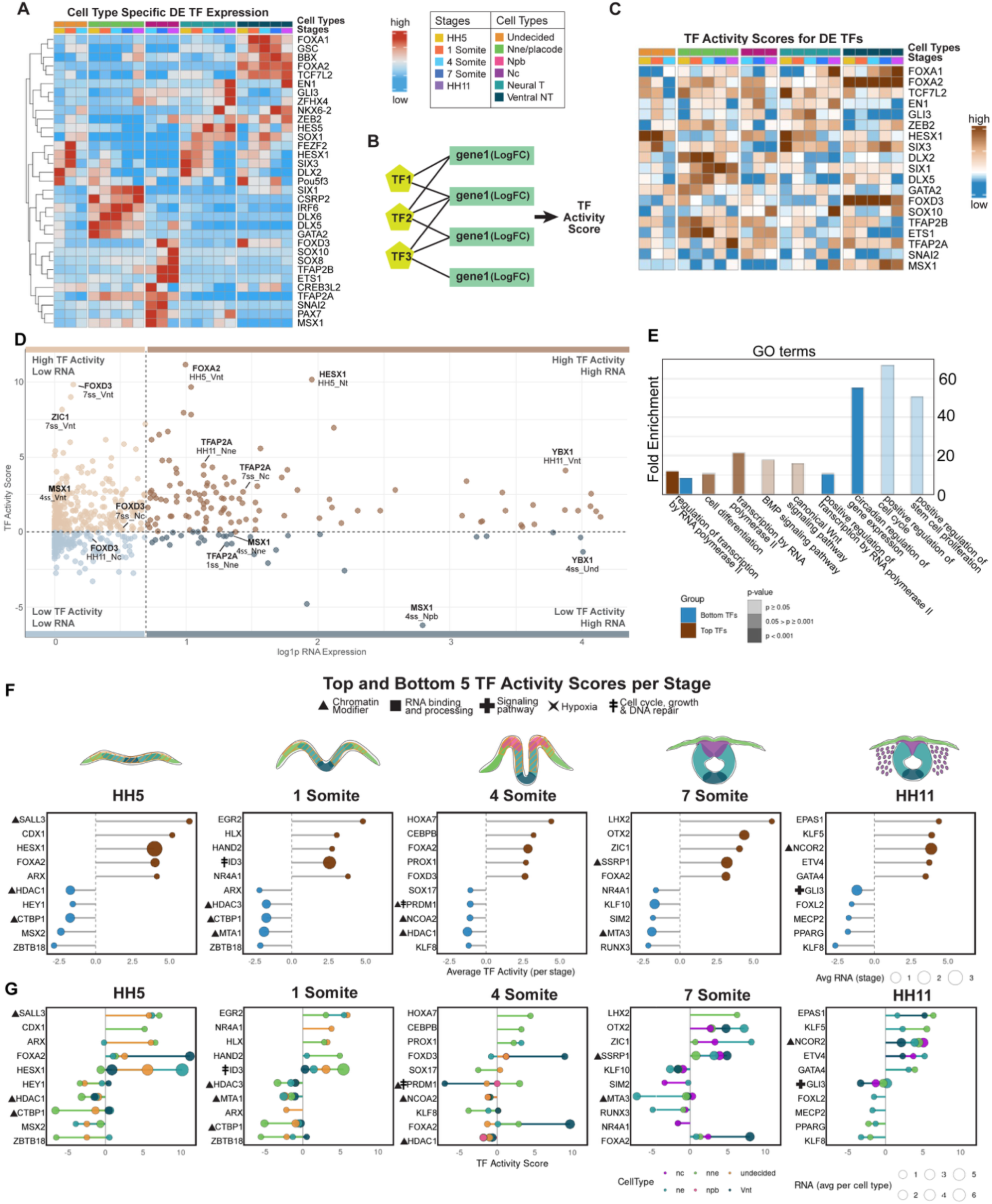
Downstream activity cannot be predicted from RNA expression levels. **A**) Heatmap presenting top differentially expressed (DE) transcription factors (TFs) in ectodermal cell types (minimum percent = 0.25, log fold-change threshold = 0.2). **B)** Schematic illustration of the decoupleR workflow, highlighting that in our analysis we used log fold change values from the DE genes identified across our 21 stage-cell type specific identities to compute TF activity scores (**Supplemental Fig. S2A**). **C**) A heatmap showing the TF activity scores of the same TFs as in panel A (Only TFs with available activity scores are displayed). **D**) Scatter plot comparing RNA expression and TF activity score. Thresholds for RNA expression (log1 value 1 to distinguish high vs low) and TF activity (score of 0 to separate positive vs negative) are used to divide TFs into four categories. Example genes that show similar transcript levels but distinct activity scores in different cell types and developmental stages are highlighted. **E**) Genes with the overall highest and lowest canonical downstream scores are plotted per developmental stage. Stem length reflects inferred activity, while bubble size represents RNA expression level, thereby integrating both measures in a single view. Bubble sizes correspond to RNA expression level. **F**) A breakdown of the genes with top and bottom activity scores per cell type. **G**) Top gene ontology terms consisting of genes with top and bottom (n= 30 each) gene activity scores at early neurulation stage (1som). Bars show fold enrichment, with significance represented as –log10(Benjamini FDR). The rest of the stages are shown in **Supplemental Fig. S2B.**

To further examine the relationship between RNA abundance and inferred TF activity, we plotted RNA expression against activity scores for each stage-specific cell types (**Fig. 2D**) and highlighted selected TFs on the scatter plot. We then categorized TFs into four conceptual groups using two thresholds: an activity threshold of 0, separating positive from negative activity, and a log1-scaled RNA expression threshold of 1, distinguishing high and low expression. Two of these followed the expected correlation between RNA and activity: high expression / high activity, representing TFs with both elevated transcript levels and strong functional influence, and low expression / low activity, corresponding to TFs with minimal involvement. In contrast, the other two categories reflected discordance between expression and activity: high expression / low activity, indicating factors that are abundantly transcribed but show limited regulatory output, potentially due to post-transcriptional or post-translational repression or non-canonical downstream activity not predicted by the algorithm; and low expression / high activity, suggestive of factors exerting strong regulatory impact despite modest transcript levels. For example, Ybx1, a multifunctional RNA/ DNA binding and stabilizing gene (51), although expressed at similarly high levels, has a high canonical activity score in the post-neurula HH11 ventral neural tube, whereas in the 4 som mid-neurula undecided cells show a negative score value, suggesting a different, non-canonical role or a repressed status. Similarly, although TfAp2a expression levels are similar in all three cell populations, the activity score is high in the premigratory, 7som neural crest cells and in the epidermis after neurulation (HH11) but negative at an earlier stage in the epidermis, reflecting a difference in the downstream activity between these stages (**Fig. 2D**). Collectively, these comparisons demonstrate that TF RNA expression alone is not always a reliable indicator of functional activity, highlighting the added value of activity-based inference for resolving regulatory states.

Next, we identified the top five highest and bottom five lowest activity scores at each developmental stage (averaged across all cell types) and visualized them using lollipop plots (**Fig. 2E**). This stage-by-stage comparison highlights TFs with consistently strong activity as well as those with minimal regulatory impact, offering insight into stage-specific regulatory landscapes. For a more detailed resolution, the same plots generated at the cell type level are provided in **Fig 2F, Supplemental Table 3).**

#### Genes with highest canonical downstream activity scores during ectoderm patterning

The results highlight a few genes per cell stage that show particularly high canonical activity, or inactivity, in the ectodermal patterning process (**Fig. 2E**). These include Sall3, which regulates differentiation of ectoderm from pluripotent stem cells (52), and has a high activity score across all ectodermal domains (**Fig. 2E, F**). Similarly, the stem cell regulator Hesx1 (44) stands out with its high expression and canonical activity levels, which mainly results from activity in the undecided pan-ectodermal stem cells and the developing neural plate after gastrulation, and ID3 activity across the entire ectoderm reflects high proliferation in the early neurula (1som). On the other hand, Forkhead box transcription factor FoxA2 is selected within the top 5 genes according to high canonical activity at post gastrula and end of neurulation stages, although the activity mainly comes from the ventral neural tube. Similarly, the homeobox gene family members of the caudal type family, Cdx1 (at post gastrula stage HH5), and the LIM family transcription factor Lhx2 (at end-of neurula stage HH9), which is known for its role in epidermal development (53), are highlighted although the activity solely stems from the developing non-neural ectoderm. The orphan nuclear receptor Nr4a1, mainly known for its function in suppressing differentiation of T-cells (54), is highlighted for highest activity in the early neurula (HH7, 1 som) with the activity stemming from the undecided pan-ectodermal stem cells. Intrestingly, the well-known Forkhead box transcription factor and neural crest specifier gene Foxd3, shows high canonical activity at mid-neurula stage in the ventral neural tube. At the end of neurulation, the homeodomain binding anterior cranial axial level determinant Otx2 shows canonical activity across all cell types. After neurulation (HH11), the transcriptional co-repressor Ncor2, known for recruiting histone deacetylases (Hdac3 in particular) as part of the SMRT repressor complex (55) as well as the hypoxia-responsive gene Epas1 (*aka* Hif2a), that plays a role in trunk neural crest development (56), showed high canonical activity across all cell types that mainly stemmed from the developing epidermis and ventral neural tube (**Fig. 2E, F**).

#### Genes with lowest downstream activity (negative canonical activity scores) during ectoderm patterning

The genes that were highlighted in the top 5 across the early ectoderm (HH5 and 1som) ectoderm for their inactivity of canonical downstream target events included histone deacetylases (Hdac) 1 and 3, Mta1 and 3, components of the nucleosome remodeling and deacetylase (NuRD) complex linked to survival and reprogramming of ESCs (57) that also have adenine methylation abilities (58), and transcriptional repressors Zbtb18 and Msx2 that have, despite being expressed in significant amounts failed to activate their canonical downstream input. Interestingly, the C-terminal binding protein Ctbp1 that recruits Hdacs and represses pro-apoptotic genes (59) showed a highly negative score in the developing epidermis, is also a co-repressor that binds to Zbtb18, which also was among the genes with high inactivity across the early ectoderm with the scores stemming from the non-neural ectoderm and the undecided pan-ectodermal stem cells. However, they showed modest but positive canonical activity in the neural tube populations in line with its reported required role in guarding neural development by repressing non-neuronal transcriptional networks (60). Msx2, on the other hand is a broadly used marker of the neural plate border at a slightly later stage (1 som/4som) that regulates the expression of genes in the neural and non-neural boundaries (61), but was highlighted across the ectoderm for its negative score right after gastrulation. Similarly, at the late neurula stages (4som, 7som), the genes that fail to induce canonical activity in all cell types include Hdac1, Klf8 and Klf10, which is known to bind Sp1 enhancer sites to repress TgfB and Smad signaling (62), as well as the hormone-receptor co-activator and epigenetic modulator Ncoa2. On the other hand, the master transcriptional repressor and histone acetylase recruiter Prdm1 (*aka* Blimp1) was included in the top 5genes with negative activity scores, which stemmed mainly from the ventral neural tube, whereas similar expression levels in the non-neural ectoderm show high, canonical downstream activity. Interestingly, Sox17, a key transcription factor of endodermal fate gene regulatory network (63), was among the highlighted genes with negative scores, which mainly stemmed from the developing non-neural ectoderm. At the end of neurula stage (7 som), Sim2, a HLH PAS family transcriptional driver of midline development (64), was highlighted with the negative activity score mainly coming from the neural crest. Nr4a1, which was highlighted for its canonical downstream activity in the undecided stem cells at 1 som stage, also showed a negative score solely sourced from the neural crest. Finally, at post-neurulation stage (H11), Gli3, the modulator of Hedgehog signaling (65) was highlighted with non-canonical or repressed signal scores in all cells except for modest positive activity in the dorsal part of the neural tube (**Figs 2E-F**).

Finally, we checked the gene-ontology terms that were associated with the top 30 genes on the overall activity list of transcription factors. After gastrulation (HH5), Regulation of Epithelial Cell Proliferation was highlighted, whereas dorsoventral axis specification stood out at mid-neurula stages (1,4 and 7som), epithelial cell development was prominent again at end of neurulation, whereas response to oxidative stress and glucose were highlighted at the post-neurulation stage (**Fig. 2G and Supplemental Fig. 2B**). Taken together, these results bring novel insight about less studied genes that may play major roles during ectodermal patterning and revealed that downstream activity of a given gene may vary between neighboring domains and developmental stage. This result prompted us to ask what are the genes that **1**) are continuously expressed and show canonical downstream activity throughout development of a given ectodermal domain **2**) are broadly expressed but show positive canonical downstream activity only in a given cell type, **3**) that fluctuate their canonical activity from high to negative values within a given cell type or developmental stage indicating different roles in different cell types?

### Core Cell type specific genes with consistent downstream activity throughout ectoderm patterning

Next, to identify potential core regulators of each ectodermal lineage, we focused on transcription factors that displayed consistent activity across all developmental stages within a given cell type (**Fig. 3A-E, Supplemental Table 4**). Specifically, we selected TFs with RNA expression levels above zero and TF activity scores that remained positive throughout neurulation in a single ectodermal domain, suggesting a continuous regulatory role over time. This filtering step was applied uniformly and is maintained across all subsequent analyses. Of the list of selected genes with continuous TF activity the genes with the highest average RNA expression levels were then highlighted as top genes in lollipop plots. In parallel, we also examined TFs that consistently exhibited negative activity scores across stages, which may indicate persistent repression of their canonical target programs or activity of a yet unknown downstream program (**Fig. 3A-E**).

**Figure 3.**
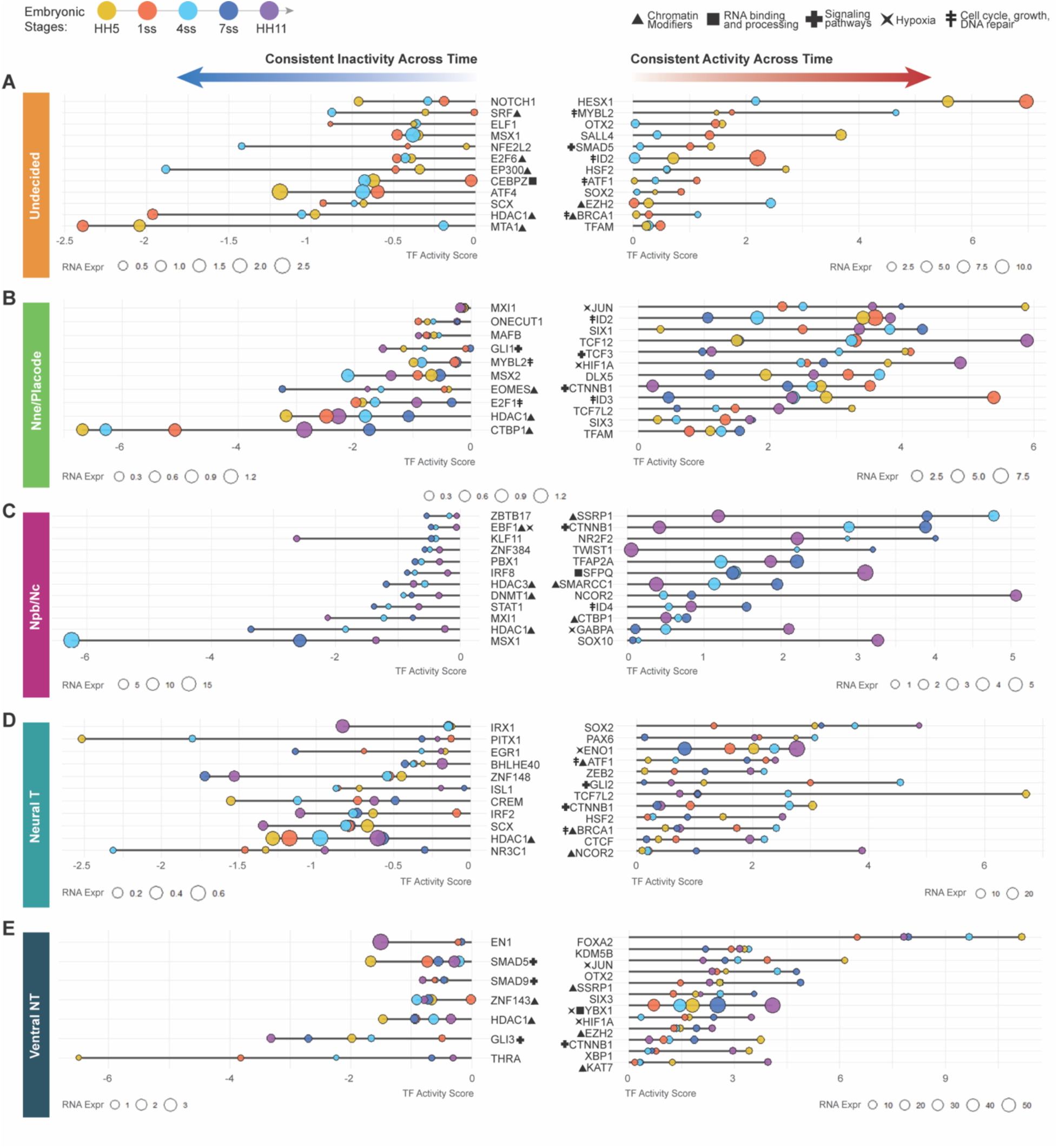
Canonical transcription factor inference analysis shows core cell fate driving genes by highlighting TFs with a consistent activity pattern in a given ectodermal cell type as shown on lollipop plots. Transcription factors displaying continuously positive or negative activity scores, throughout developmental stages of **A)** undecided pan-ectodermal stem cells, **B**) non-neural ectoderm and placodal cells, **C**) neural plate border and neural crest cells, **D**) dorsal neural plate and tube cells and **E**) ventral neural plate and tube cells. For this analysis TFs with RNA expression values below zero were excluded. Bubble sizes correspond to RNA expression levels and are shown relative to the scale used within each individual plot. Stem length reflects inferred activity.

#### Highest continued canonical downstream activity in undecided pan-ectodermal stem cells

In the undecided pan-ectodermal cells, the results show continuous activity of the known stem cell genes HesX1 (44), as well as Sall4 and Sox2, which are part of the pluripotency complex (66), although Sox2 is also a main driver of the neuronal fate in neural progenitors in the neural plate (67). Also, cell cycle and cell growth activators Id2 (68), Mybl2, and Atf1, and genomic stability maintaining Brca1, multifunctional heat shock protein Hsf2, and Ezh2, histone methyltransferase that is a part of the repressive polycomb protein complex 2 (69) showed continuous activity. Smad5, a downstream effector of BMP-signaling, a major driver of non-neural ectoderm, was also highlighted. In sum, the genes that displayed consistent canonical activity in the stem cell population have reported roles in promotion of stem cell maintenance, neural and non-neural fate as well as cell growth (**Fig 3A**).

#### Lowest continued canonical downstream activity (negative score) in undecided pan-ectodermal stem cells

Conversely, the genes that are expressed and repressed, or alternatively activate non-canonical downstream activity, included the cellular stress responder Atf4, Mta1, essential for ESC epigenetics and survival (57), Srf, which associates with Sox2 and Nanog and contributes to the formation of long range chromatin loops to create 3D pluripotency hubs (70), the nucleolar gene Cebpz, essential for rRNA processing and maturation (71), E2f6, a member of the polycomb repressive complex that initiates stable epigenetic silencing via promoter methylation during embryogenesis (72), the histone acetyltransferase EP300, the histone deacetylase and transcriptional silencer Hdac1, and Elf1, a hematopoietic Ets family TF not linked to ectoderm before showed consistent negative activity scores. The known neural plate border repressor gene Msx1 (61), as well as the neuronal fate determinator Notch1, which is also implicated in differentiation of multiple other tissue types were also highlighted (**Fig 3A**).

#### Highest continued canonical downstream activity in non-neural ectoderm

The development of the non-neural ectoderm at the midbrain level is shaped by continuous canonical downstream activity of the known epidermal fate determinator Dlx5 (73) and the placodal gene Six1 (74). Additionally, cell cycle regulators Id2, Id3 and the E-box binding transcriptional regulator Tcf12 that is associated with multiple downstream activities as well as non-canonical Wnt-signaling (75). Interestingly, other canonical Wnt-signaling regulators Ctnnb1 (*aka* Beta-Catenin), the Lef-repressor Tcf3 (76) as well as the key mitochondrial DNA transcription factor Tfam that activate stem cell genes downstream of Wnt-signaling (77) and the hypoxia induced Hif1a, which both are active throughout neurulation stages in the non-neural ectoderm. Finally, the proto-oncogene and Ap-I family gene Jun (*aka* C-Jun), which also is functionally connected to embryogenesis and cellular protection of oxidative stress (78) was also highlighted (**Fig. 3B**).

#### Lowest continued canonical downstream activity (negative score) in non-neural ectoderm

Conversely, the genes that showed top negative activity scores, in addition to the genes listed already in the overall top genes of negative scores (Msx1/2, Ctpb1, and Hdac1), consisted of Mybl2, which showed consistent canonical activity in the undecided population (**Fig. 3A**), suggesting separate roles in the neighboring cell types. Additionally, the cell cycle regulator E2F1, Mafb, Eomesodermin (Eomes), that plays a role in meso- and endoderm lineage specification by driving epigenetic changes (79) also shown to be activated by Retinoic acid (RA) signaling (80) and Onecut (81), which regulate development of multiple tissues although not reported in skin formation, as well as Gli1, the dedicated activator of the Shh-signaling pathway (82) showed negative activity, although their expression levels were notably low. In sum, Wnt-signaling related genes were highly active throughout non-neural ectoderm development. Chromatin modifiers, and the known neural plate border repressors Msx1/2 as well as the dorsoventral polarity determining Shh downstream activator Gli1 were amongst the repressed/non-canonically functioning genes to shape the formation of the epidermis, and Mybl2 showed a different downstream activity as in the undecided cells (**Fig. 3B**).

#### Highest continued canonical downstream activity in neural crest

The genes that showed highest canonical activity in the developing neural crest cells (neural plate border and specified neural crest) starting from mid-neurula stage (4ss) until migratory stage (HH11) included Tfap2A, which was highly expressed and active throughout the specification and EMT process, confirming its known role during neural crest specification (83). Also, the known neural crest marker Sox10 was continuously active, with highest activity and expression found in the migratory cells, as well as Twist1 and Ctnnb1 (B-catenin), important for EMT in the remodeling of adherence junctions also independent of Wnt signals (84). Notably, despite Ctnnb1, other Wnt signaling activity was not highlighted. Also, another cell cycle activator member of the inhibitors of DNA binding protein family, ID4, was active throughout the stages. The analysis also revealed epigenetic modifier genes for neural crest formation like Ssrp1, the nucleosome reorganizer and component of FACT (85), and the component of the SMRT co-repressor complex Ncor2 that recruits histone de-acetylases (55), which was previously identified to be downregulated in murine neural crest cultures in response to RA signaling (80), and the nuclear receptor Nr2f2, which regulates neural crest transition to ectomesenchyme in zebrafish and is involved in melanoma metastasis (86, 87). Additionally, Sfpq, which is a RNA splicing protein the loss of which is shown to cause CNS and neural crest defects in zebrafish (88) and Smarcc1 (aka Baf155, Srg3), a core component of the SWI/SNF chromatin remodeling complex, depletion of which is linked to hydrocephalus (89), was highly expressed and canonically active through all stages of neural crest development. Finally, in line with the maintenance of pluripotency gene activity being part of neural crest lineage establishment (22), Gabpa (*aka* Nrf2) a mitochondrial biogenesis gene that regulates cellular defense against toxic and oxidative insults and and newly identified as a master regulator of epiblast stage transition from the inner cell mass before gastrulation (90, 91) showed high expression and activity in the neural crest lineage (**Fig. 3C**).

#### Lowest continued canonical downstream activity (negative score) in the neural crest

On the other hand, Msx1, shown to shape the specification of neural plate border to form neural crest and placodes (92) stood out of the genes highlighted as repressed or with non-canonical activity. The top list also included two cMyc regulating genes: Mxi1, a Max-binding tumor supressor that antagonizes cMyc among other things (93, 94) and Ztbt17 (*aka* Miz-1) which we have previously shown to regulate the neural crest stem cell pool via binding to c-Myc via a non-canonical pathway (95, 96). Additionally HDAC 1 and 3 were again highlighted as well as the DNA methyltransferase Dnmt1, of which the canonical role is to maintain DNA methylation patterns in daughter cells and is important for later neural crest derived functions in orofacial clefting morphogenesis (97), Pbx1p that has been shown a role in Xenopus neural crest development in collaboration with Meis1 (98), palate morphogenesis (99), and NC derived adipocyte formation (100). Finally, amongst the highlighted genes was Ebf1 transcription factor and oncogene that is shown to assemble a transcriptional complex with Hif1a to suppress p300 activity in triple negative breast cancer (101) and is included in the neural crest gene regulatory network (2). Additionally novel genes like IRF8, known for its role in hematopoietic lineage differentiation, Stat1, known for its role in innate and adaptive immunity, Znf384, which promotes EMT in breast cancer cells and is linked to Zeb1 (102), and the Krueppel-like factor Klf11, known for its role in extracellular matrix regulation by repression of Collagen 1a2 (103) and involvement in glucose metabolism (104) were highlighted. It remains unclear whether these genes are actively repressed in the developing neural crest or whether they activate yet unknown downstream activity networks in these cells (**Fig. 3C**).

#### Highest continued canonical downstream activity in neural plate/neural tube (excluding the ventral NT)

Next, we analyzed the genes highlighted based on their top activity in the developing dorsal and lateral parts of the neural tube (**Fig. 3D**), which contained the known early neural fate determining gene Sox2, Pax6 (36), (105) and Zeb2 (106), which is also essential for finalizing neural crest EMT (107). Unexpectedly, high activity of Gli2, the downstream activator of the ventraling Shh signaling is shown in the dorsal neural tube (108). High activity of the HDAC complex member Ncor2 (*aka* Smrt)(55), especially in the post-neurulation stage was also shared with the neural crest. Similarly, activity of the growth promoting and chromatin modifier recruiter genes Brca1 (109) and Atf1 (110) was shared with the undecided pan-ectodermal stem cells (Fig 3A,C). Interestingly, Eno1, the enolase enzyme of the glycolysis pathway, which can also produce the transcription factor c-Myc promoter binding protein-1 (Mbp-1) via an alternative translation initiation mechanism (111) is highly expressed and active in the neural tube cells. Also, the three-dimensional chromatin enhancer insulator protein Ctcf that regulates gene regulatory networks in multiple differentiating cells, including the brain during development (112) is highly active throughout neural tube formation. Finally, similar to the non-neural ectoderm Tcf7l2 (*aka* Tcf4) and its binding partner CTNNB1 (*aka* Beta-Catenin) that together activate signaling downstream of Wnt are highly expressed and active in the neural tube (**Fig. 3D**).

#### Lowest continued canonical downstream activity (negative score) in neural plate/neural tube (excluding the midline ventral NT)

Conversely, the genes in the plot that highlights continues negative downstream activity across neural tube development were all expressed in very low levels. They included Hdac1, Irf2, which is known for its role in development of natural killer immune cells, the connective tissue related gene, Scx, the mesodermal gene Pitx1, Znf148 that is broadly involved in embryogenesis, and Crem that can act both as an activator or repressor of cAMP dependent signaling (113) and glucocorticoid receptor gene Nr3c1.

#### Highest continued canonical downstream activity in ventral neural plate/tube

Finally in the ventral neural tube, of all the highlighted genes, Foxa2 activity was maintained the highest throughout neurulation, consistent with its role in floorplate activation induced by Yap signaling mediated tissue stiffness (114). Interestingly, four of the top activity genes Jun (78), Hif1A, Six3 and Ctnnb1, are shared with the developing non-neural ectoderm (**Fig. 3B and E**). Similarly, the polycomb gene Ezh2 that catalyzes the methylation of H3 histone of H3K27Me3 (69) and the anterior body axis determining Otx2 are shared with the undecided cells (**Fig. 3A and F**). Ybx1 is a multifunctional RNA/DNA binding protein with broad roles in development and cancer (51) but its role in ventral neural tube, floor plate or motor neuron development remains unknown. Our data shows that it’s abundantly expressed and highly active in the ventral neural tube, and it also has been shown to bind to Hif1a mRNA in hepatocellular carcinoma cells (115). Histone demethylase Kmd5b (*aka* Jarid1B), which is known for its role in inhibiting differentiation also in neural stem cells (116), Xbp1, a cellular stress response associated transcription factor that has been shown to also enhance hif1a target gene expression in aggressive breast cancer (117), Ssrp1, a part of the Histone chaperone and FACT unit (85), and the histone acetyltransferase Kat7 (*aka* HBO1), which was recently shown to function as a H3K14 acetylase that is indispensable for re-activation of repressed genes to maintain stem cell plasticity in neural stem cells (118)(**Fig. 3E**).

#### Lowest continued canonical downstream activity (negative score) in ventral neural plate/neural tube

The genes with the negative activity score in the ventral neural tube included Gli3, an activator of Shh signaling and a repressor of it in the absence of Shh activity (119), En1, a regulator of wnt signaling induced neurogenesis the midbrain (120), Hdac1, the thyroid hormone receptor ThrA, that maintains the neural progenitor pool in the forebrain (121), and the zinc finger protein Znf143, which mediates formation of CTCF-bound promoter-enhancer loops in hematopoietic stem cells (122). Furthermore, the Bmp and Tgf-beta signaling pathway inhibitors Smad5 and Smad9 are among the top genes that show negative or non-canonical activity in the ventral neural tube (**Fig. 3E**). Interestingly, Smad5 was highlighted among the top genes with canonical activity in the undecided pan-ectodermal stem cells, suggesting a different role in the ventral neural tube (**Fig. 3A and E**).

Taken together, the data highlights novel and understudied transcription and epigenetic factors, as well as downstream effectors of signaling pathways with high canonical and, on the other hand, non-canonical or repressed activity that shape the cell type specific domains during ectodermal patterning by showing distinct activity in the domain throughout the process, suggesting a core function for the cell type. Interestingly, the data shows non-canonical or repressed activity for the histone deacetylase 1 (Hdac1) in all cell types, and genes that were shared between several cell types such as Ctnnb1 (beta-catenin), which was highly active in all cell types except the undecided stem cells, suggesting high adherence junction modifications and/or Wnt signaling activity response across cell types during ectodermal patterning. Lastly, the data highlighted genes that were canonically active in one cell type but non-canonical/ repressed in another cell type, such as Ctbp1 (neural crest vs nonneural ectoderm) and Smad5 (stem cell vs ventral neural tube), suggesting different downstream activity patterns in different ectodermal cells. All cell types highlighted known genes in addition to the novel ones, increasing the credibility of the data, thus provides new candidates for further investigation and for regenerative purposes (**Fig. 3A-E**).

### Domain-Specific Downstream Activity of Transcription Factors

Developmental biology has for decades heavily relied on *in situ hybridization,* more recently complemented with RNA sequencing, to address importance of genes for a given cell type during organogenesis. Epigenetic regulators are often broadly expressed across cell types and transcription factors, although known to bring lineage specificity, are rarely, if ever, expressed solely in one cell type, as also shown in our RNAseq data set (**Fig. 1A and Supplemental Fig. 2A**), and co-binding to different factors in different gene regulatory networks is known to broaden their repertoire of executing different tasks. The fact that Fig 3 highlighted genes that were equally expressed in two cell types but showed positive vs negative activity in them prompted us to investigate, whether we could differentiate between the function of transcribed genes in neighboring cells based on their downstream activity by dissecting the information from scRNAseq data alone. Previous studies also indicate that co-expression of several gene regulatory networks of neighboring domains define an uncommitted status of fate determination (22, 24), but the downstream activity of the genes has not been investigated in these studies. Thus, we hypothesized that the embryo may rely on differential TF activity, rather than expression levels alone, to sharpen cell type boundaries and reinforce lineage specification. To test this idea, we searched for TFs with distinct activity profiles at each developmental stage. Specifically, we classified TFs as 1) exclusively active when canonical activity was positive (>0) in one cell type but negative (≤0) in adjacent cell types, and 2) as gradient active when high activity (exceeded 1) in one cell type while remaining intermediate (0–1) in neighbors. We then highlighted the top TFs within each category and summarized them across stages in a composite schematic (**Fig. 4A-E, Supplemental Tables 5-9**).

**Figure 4.**
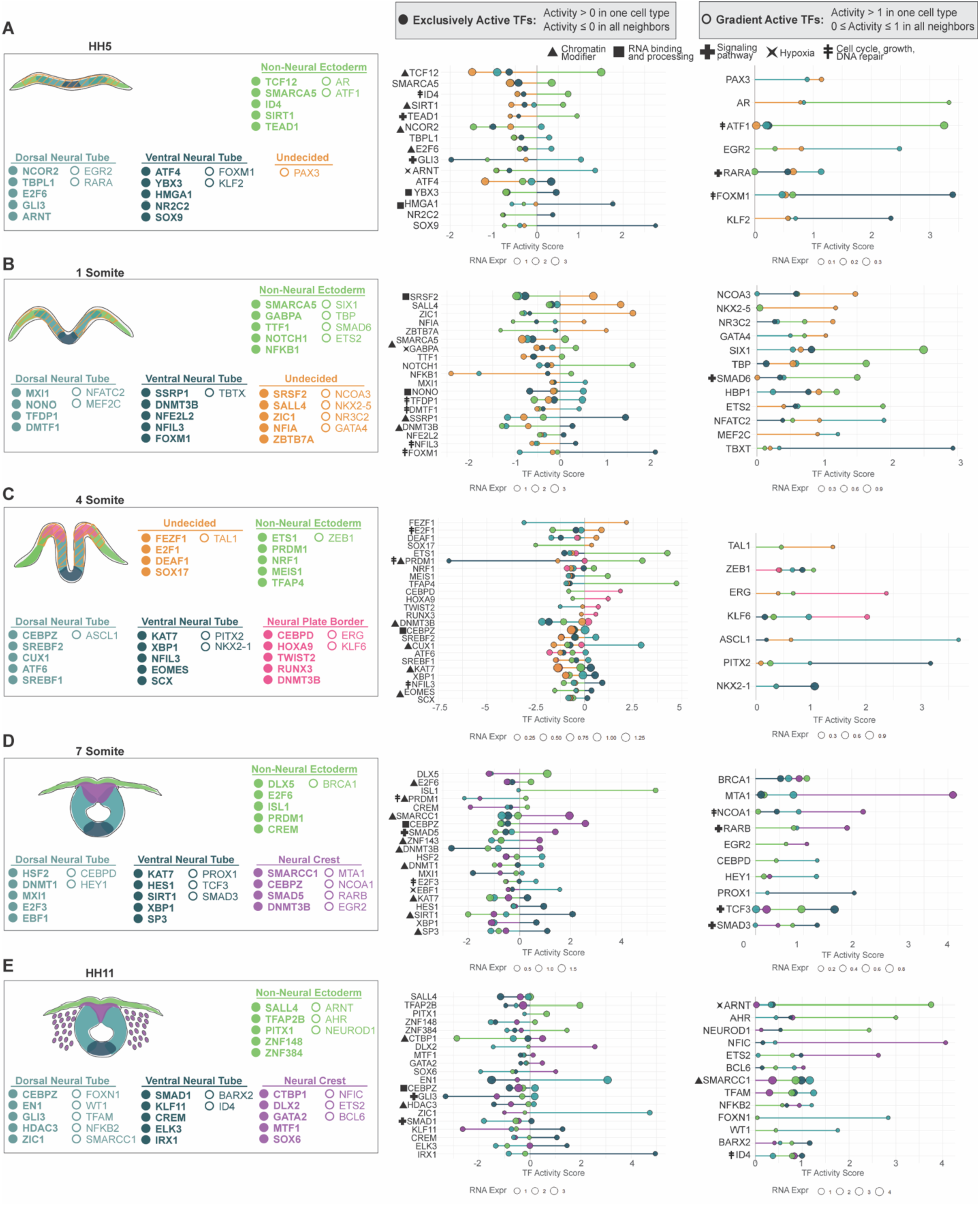
Canonical transcription factor activity profiles reveal cell-type specific exclusive and gradient activity patterns across developmental stages. Lollipop plots showing transcription factor activity patterns across ectodermal cell types at HH5, **B**) HH7 (1 som), **C**) HH8 (4 som), **D**) HH9 (7 som) and **E**) HH11. For this analysis TFs with RNA expression values below zero were excluded. Bubble sizes correspond to RNA expression levels and are shown relative to the scale used within each individual plot. Stem length reflects inferred activity.

#### Highest canonical downstream activity after gastrulation (HH5)

##### Undecided pan-ectodermal stem cells

The results show that right after gastrulation, in line with the hypothesis that the newly developing ectoderm is very plastic and expresses low levels of multiple ectodermal genes (22), the undecided cells didn’t contain genes that would be uniquely active only in this cell population. The neural plate border gene Pax3 was highlighted with slightly increased activity as compared to the developing neural plate (neural tube), although the RNA expression levels in the gradient activity group for HH5 were low (**Fig. 4A**).

##### Non-neural ectoderm

On the other hand, signs of specification in terms of unique activity in only a given cell type were detected in both neural and non-neural ectoderm at this early developmental stage. Specifically, activity of Tcf12, associated with non-canonical Wnt signalling, and cyclin D regulation (75), the cell cycle inducer Id4, as well as Tead1, the cell growth promoting downstream transcription factor of Hippo signaling pathway but also independent of it (123) and the multifunctional NAD+ dependent histone deacetylase Sirt1 that is known to modulate neuronal vs glial differentiation via Notch signaling regulation (124) were among the genes with unique activity in the non-neural ectoderm (**Fig. 4A**).

##### Neural plate (excluding midline / ventral neural plate)

In the early developing neural plate that excludes the midline, on the other hand, unique activity of the deacetylase recruiter Ncor2 (55) and Tbpl1, that among other targets represses cell growth via transcriptional activation of p53, p63 and p21 (125), cell cycle repressor and recruiter of the Bmi1 containing polycomb repressor complex E2f6 (126), Gli3 that inhibits Shh signaling pathway in the absence of the ligand (119), as well as the Aryl hydrocarbon receptor Arnt (aka Hif1b) that senses hypoxia and pollutants such as dioxin and dimerizes with Hif1a (127, 128) were detected (**Fig. 4A**). Furthermore, the Retinoic acid receptor alpha Rara activity was distinguished, in line with studies on RA-signaling regulating the duration of Bmp signaling via promoting degradation of phosphorylated Smad1 in the neural tube (129), although the RNA expression level was extremely low (**Fig. 4A**).

##### Ventral neural plate

Finally, right after gastrulation, genes with canonical activity solely in the developing ventral neural tube in the middle of the neural plate included Atf4, a redox-regulated cell death promoting transcriptional activator of the CREB family (130), the RNA binding translation modulator Ybx3, the loss of which causes infertility (131), as well as, although expressed in low level, the neural crest and chondrocyte lineage activator Sox9, which is also known for its role in maintenance of the notochord (132) that forms underneath the ventral neural tube, as well as the non-histone protein Hmga1 that modifies DNA structure (133)(**Fig. 4A**).

#### Highest canonical downstream activity at early neurula stage (HH7, 1 som)

##### Undecided pan-ectodermal stem cells

In the early neurula at stage 7 (1som), the genes in the undecided pan-ectodermal stem cells at midbrain level of the chick embryo that showed high activity in only one cell type included Srsf2, which controls RNA splicing gene expression via multiple regulatory ways (134), the pluripotency gene Sall4 (66), and the neural plate and neural plate border inducing transcription factor Zic1 that regulates early neural crest and neural induction (135, 136). The multifunctional oncogenic driver and transcriptional repressor Ztbt7a (137), and the Nuclear factor 1 family transcription factor activator Nfia, the mutation of which causes the NFIA-related neurodevelopmental disorder that also presents with craniofacial defects (138)(**Fig. 4B**).

##### Non-neural ectoderm

The genes that show canonical activity only in the non-neural ectoderm at early neurula stage included the notch-signaling receptor Notch1 that activates Hes transcriptional repressors during cellular fate decision and maintain the cell in a progenitor stage (139), the Imitation Switch (ISWI) gene Smarca5 that encodes the ATP-dependent chromatin remodeling protein Snf2h and is known to regulate cell differentiation (140), the master regulator of pluripotency GabpA (90) as well as NfkB1, which in addition to its immunological role is ubiquitously expressed in all cell types in the embryo except the embryonic stem cells and plays a role in neuro-ectodermal and mesodermal development (141). Additionally, the Bmp inhibitor Smad6 associated with neurocristopathies (142) and the cranial placodal driver Six1 as well as the transcription factor Ets1, which is associated in development of multiple tissues, were highly canonically activate in the non-neural ectoderm during early neurulation although low activity (< score value 1) was detected in the other ectodermal domains as well (**Fig. 4B**).

##### Neural Plate (excluding ventral midline)

Next, we analyzed the canonically active genes unique for the neural plate. These included Mxi1, a bHLHzip transcription factor that competes with Max to bind to Myc to repress its downstream activity and is shown to promote early neural fate in the neurula (94). Furthermore, other high activity genes included Nono, a X-linked multifunctional nuclear protein involved in RNA splicing, DNA repair and transcriptional regulation mutations of which cause neurodevelopmental and cardiac defects (143), Tfdp1, which dimerizes with the cell cycle activator E2f to enhance its activity (144), as well as the Cyclin D-binding Myb-like transcription factor 1 Dmtf1, which, on the other hand, binds to the Sarf promoter to activate the apoptotic p53 pathway (145)(**Fig. 4B**).

##### Ventral neural plate (midline)

Finally, the genes highlighted for their unique canonical activity in the ventral neural tube at early neurulation stage included Ssrp1, which is a subunit of the chromatin remodeling FACT complex (85) and Foxm1, which controls a network of microRNAs that are required for neural stem cell self-renewal in a Shh and Nanog driven fashion (146), and which was also distinguished by its activity at earlier the stage (HH5, **Fig. 4A**) in the ventral neural plate. Furthermore, unique but lower activity levels were highlighted for the DNA methyl transferase Dnmt3B which is a major chromatin regulator for gene silencing and associated with inhibition of premature neuronal differentiation (147), neurodevelopmental defects (148) as well as for methylation of the Sox10 promoter after neural crest emigration (149). Nfe2l2, that encodes Nrf2, a master regulator of defense against oxidative stress in mammalian cells that is linked to neurodevelopmental development (150), and the repressive transcription factor Nfil3 which binds RNA-polymerase II and enhances cell growth and survival of (ventral neural tube derived) motorneurons) and has shown antagonistic action with other Par protein family members (151–153) showed high downstream activity. Furthermore, although expressed in extremely low level, Tbxt, the master regulator of mesodermal fate via enabling EMT (154, 155) was significantly high in the ventral neural tube (**Fig. 4B**).

#### Highest canonical downstream activity at mid-neurula stage (HH8, 4 som)

##### Undecided pan-ectodermal stem cells

Next, we analyzed the genes that were highlighted by their unique activity at the mid-neurula stage (4som) despite being expressed at similar levels in all ectodermal domains. In general, all the highlighted genes were expressed in low levels. The undecided pan-ectodermal cells displayed high activity of the anterior neuroectodermal zinc finger protein Fezf1, which plays a role in forebrain development (156), the cell cycle activator E2f1, and to a lesser extent Deaf1, which is important for the closure of the neuropore (157) as well as the endoderm inducer gene Sox17 that represses ectodermal fate via Wnt signaling (63) (**Fig. 4C**).

##### Non-neural ectoderm

The genes highlighted in the non-neural ectoderm consisted of Prdm1 (*aka* Blimp1) that promotes cell growth by repressing p53 (158) and is mostly known for its role in immune and germ cells, and surprisingly Ets1, which is a part of the neural crest gene regulatory network and required for its migration and shown to bind to Hdac1 to attenuate BMP signaling in frog neural crest (159). Finally, the cell cycle progression promoting oncogene Tfap4, which among other genes acts downstream of Myc and N-Myc in activating EMT and proliferation and is associated with stemness control (160) as well as Meis1, which is involved in epidermal stem cell and skin tumorigenesis regulation (161) showed high and unique activity in the developing skin at mid-neurulation stage (**Fig. 4C**).

##### Neural plate border (future neural crest)

All the genes highlighted for their canonical downstream activity in the developing neural crest domain at the neural plate border region were expressed at very low level. Of these, the highest activity was detected by Cebpd, a transcription factor that regulates multiple genes related to hypoxia response in Glioblastoma and other cancers and developing organs, and has been linked to ECM modifications via Egfr/PI3K signaling (162) and the DNA methyltransferase Dnmt3b (148) showed moderate activity but low expression similar to Runx3, HoxA9 and Twist2 (**Fig. 4C**).

##### Neural Plate, neural folds (excluding ventral midline domain)

In the rising neural folds of the developing dorsal neural tube, the pioneer factor and chromatin structure regulator Cux1 was the only strongly highlighted gene. It is target of Tgf-beta signaling (163) and associated with several cancers that has also been linked to neural stem cell fate during corticogenesis (164)(**Fig. 4C**).

##### Ventral neural plate/tube

In the ventral portion of the neural folds/tube, Nfil3 (aka E4pb4), which was highlighted also at the previous stage in Fig 4B (151–153), showed the highest canonical downstream activity. Furthermore, the high activity genes included the histone lysine acetyltransferase of H3K14, Kat7 (*aka* Hbo1), a promoter of hematopoietic stem cell quiescence and self-renewal that is indispensable for post gastrulation development (165) and neural stem cell maintenance (118). Kat 7 is also known to acetylate Ybx1 (166) which itself was also highlighted for unique activity. Ybx1 is an endoplasmic reticulum gene associated with RNA splicing, which overlaps with PRC2 distribution and inhibits H3K27me3 levels during embryonic brain lineage specification and self-renewal vs differentiation choices (167) (**Fig. 4C**). Both Kat7 and Ybx1 were highlighted previously as genes that are active throughout the ventral neural tube development in Fig 3E.

#### Highest canonical downstream activity at end of neurulation stage (HH9, 7 som)

##### Non-neural ectoderm

Next, we analyzed the genes that were highlighted by their unique activity at the end of neurulation stage (Fig 4D). In the developing skin, Islet-1, a gene known for its role in multiple cranial placodes (168) as well as the homeobox transcription factor Dlx5, which is a known marker required for epidermal development (169) were among the genes with highest canonical activity. Furthermore, Prdm1 (Blimp1)(158) continues to have a unique activity, although lower than at the previous HH8 stage, as well as the regulator of cAMP dependent signaling Crem (113), which also showed continuous noncanonical or repressed signaling throughout neural tube development in figure 3D, were highlighted as genes with unique canonical activity in the non-neural ectoderm (**Fig. 4D**).

##### Premigratory Neural crest

The genes that show unique canonical activity in the premigratory neural crest cells included three epigenetic modifiers: Smarcc1, a core component of the SWI/SNF chromatin remodeling complex (89), DNA methyltransferase and DNMT3b (148), which showed unique activity also at the earlier neural plate border stage (HH8), Znf143, which mediates CTCF-bound promoter -enhancer loops (122). Furthermore, Cebpz (*aka* Noc1) that is essential for ribosomal RNA processing (71), and SMAD5, which is a mediator of Bmp signaling but may also function in a separate role (170), showed unique activity in the neural crest. Furthermore metastasis associated protein 1 (MTA1) that is a component of the NuRD chromatin remodeling complex and can also act as a adenine methylase and is strongly linked to EMT in various cancers (57, 58, 171) was highly active in the neural crest with only small activity in the other domains, as well as the cancer associated nuclear receptor co-activator 1 (Ncoa1) which recruits histone deacetylases and enhances expression of several downstream target genes like hif1a and vegfa in breast cancer (172, 173) and the Retinoic acid receptor beta (Rarb), mutations of which have been linked to enteric neural crest migration defects in disease (174)(**Fig. 4D**).

##### Dorsal and dorsolateral neural tube

The genes with unique canonical activity in the dorsal parts of the neural tube included the multifunctional heat shock protein Hsf2, which was continuously active throught the early stages in the undecided pan-ectodermal stem cells in fig 3A, and which has been linked to neural plate induction before (175) and the DNA methyltransferase Dnmt1, which has been shown to promote neural and inhibit glial fate choices in vitro (176). Furthermore, although expressed in low levels, the cell cycle activator E2f3, which has a reported role in regulation of cell cycle exit upon neural stem cell differentiation (177) and Ebf1, which plays a role in neuronal differentiation of telencephalic medium spiny neurons (178) show unique activity in the dorsal portion of the closed neural tube. Finally, the non-oscillatory Notch effector gene Hey1, which is key for the transitioning of embryonic neural stem cells into quiescent adult stem cells of the subventricular zone (179) was highlighted as a gene with significantly higher activity in the dorsal neural tube as compared to neighboring domains (**Fig. 4D**).

##### Ventral neural tube

In the ventral part of the neural tube, genes with highest unique canonical activity consisted of the histone acetyltransferase Kat7 required for H3K14 acetylation to maintain neural stem cell plasticity (118) that already was highlighted at the previous stage in the ventral neural tube in Fig 4C and Hes1, the known effector downstream of Notch signaling that regulates neurogenesis (180). Other hioghlighted genes included The class III histone deacetylase Sirt1, which operates in an NAD+-dependent manner and is known for various cellular functions including DNA repair, cell survival, and inflammation in multiple tissues and diseases including embryonic stem cells (181), and Xbp1, which was uniquely active in the domain already at the previous developmental stage in Fig. 4C and continuously active in the developing ventral neural tube (fig 3E), which binds to DNA and RNA and enhances transcription and translation of multiple genes including enhancement of expression of Hif1a target genes in triple negative breast cancer (117, 182). Interestingly, Hif1a is one of the continuously active genes in the ventral neural tube (Fig. 3E). Also Sp3, a ubiquitously expressed transcription factor and chromatin modifier required for multiple developing organs and cancer (183), Tcf3, which acts downstream of Wnt-beta catenin in the neural tube (76), Smad3, the TgfB-Activin signaling mediator that promotes ventral interneuron fate specification and differentiation (184), as well as Prox1, a repressor which has been shown to suppress Olig2 required for proper motor neuron development in the ventral neural tube (185) were highlighted due to higher activity in the ventral neural tube as compared to other domains (**Fig. 4D**).

#### Highest canonical downstream activity at post neurulation stage (HH11, 13-14som)

##### Non-neural Ectoderm

Finally, we analyzed the transcription factors and epigenetic modifiers with highest canonical activity after neurulation has ended. At this stage the dorsal neural tube is reconstructed to form the roof plate, the neural crest cells are in the middle of their migration, and the epidermis and cranial placodes are fully specified and segregated. Interestingly, the neural crest specifier gene Tfap2b (83) that is uniformly and highest expressed in the neural crest, shows unique canonical activity in the non-neural ectoderm instead. The oral ectoderm gene Pitx1, required for the formation of the pituitary gland (186)as well as NeuroD, which is expressed in the developing cranial sensory ganglia (187) showed distinguished canonical activity in the non-neural ectoderm. Also the zinc finger genes Znf384, which are associated with cancer EMT and metastasis and Zeb1 transactivation (102) were highlighted. Furthermore, Aryl hydrocarbon nuclear translocator (Arnt *aka* Hif1b), which heterodimerizes with Hif1a to allow its stabilization and binding to DNA (128) in line with Hif1a itself being highlighted as one of the continuously active genes of the developing non-neural ectoderm in figure 3B; as well as the environmental toxin sensor gene Aryl hydrocarbon receptor AHR that interacts with Arnt for its transcription factor role, and which has been shown to activate epidermal TFAP2A transcription (188–190) were significantly higher activated in the developing skin as compared to the other cell types (**Fig. 4E**).

##### Migratory Neural Crest

Next, we analyzed the canonical activity unique for the migratory neural crest cells, which highlighted genes that are involved in differentiation of neural crest derived cell types. These included Gata2, which is expressed in neural crest derived developing sympathetic ganglia (191), the pro-chondrogenesis and anti-gliogenesis reported SoxD transcription factor Sox6 (192), which may also enhance melanogenesis (193). The HDAC recruiter and pro-apoptotic gene repressor Ctbp1 (59) also showed unique but modest activity, whereas the canonical activity of the neural crest migration and specification gene Ets1 (194) as well as Nfic, which is a key regulator of neural crest derived odontoblast formation during toot development (195), were both highlighted due to distinguished high activity as compared to other cell types (**Fig. 4E**).

##### Dorsolateral neural tube and roof plate

Next, we analyzed the genes with unique canonical activity in the reconstructed dorsal neural tube after the emigration of neural crest cells. Two well established genes of cranial neurogenesis, En1 (120) and Zic1(36, 136), showed the highest unique canonical activity, whereas low but unique activity was shown by Cebpz, which controls rRNA maturation (196), Gli3, a modulator of Shh signaling and a regulator of dorsoventral patterning of the neural tube (108) and the histone deacetylase Hdac3. Furthermore, the epigenetic modifier Smarcc1, which is linked to hydrocephalus (89), Foxn1, which is linked to neural tube defect and anencephaly (197) as well as Wt1, a marker of the radial glial (neural progenitor) cells (198) showed unique activity in the reconstructed dorsal neural tube after neural crest departure (**Fig. 4E**).

##### Ventral neural tube

Finally, the genes that showed unique high canonical activity in the ventral region of the neural tube consisted of Irx1, which is linked to dorsoventral patterning of the central nervous system (199). Lower but unique activity was also revealed for the regulator of cAMP dependent signaling Crem (113), not linked to neurogenesis before but which also showed continuous noncanonical or repressed signaling throughout dorsal neural tube development in figure 3D, and unique activity in the developing non-neural ectoderm at the previous stage in figure 4D. Klf11, which has been reported for its role as an inducer of oligodendroglial cell death downstream of Tgfß signaling (200) and Elk3, which is essential for neural crest development (201) and associated with cancer progression were also highlighted. Finally, the myogenic gene Barx2 that has been reported respond to Shh signaling also showed highest activity (202) in the ventral neural tube as compared to other cell types (**Fig. 4E**).

Taken together, the TF Activity Inference analysis revealed multiple ubiquitously expressed chromatin and histone modifiers, RNA processing genes and transcription factors, that have not been explored in ectodermal patterning before. Differences in downstream activity highlighted how individual cell types have unique activity of genes that are expressed in all neighboring domains as well, providing an important means to distinguishing functionality of transcribed genes based on RNA expression data and a large resource of knowledge for future studies on how ectodermal cell types develop (**Fig. 4A-E**).

### Downstream activity of previously known genes involved in ectodermal patterning

#### Downstream activity of previously known genes after gastrulation HH5

Finally, we wanted to analyze the downstream activity scores of genes that are commonly used as “typical markers” of the respective developing ectodermal domains. At the earliest stage after gastrulation (HH5), the pluripotency genes Sall4, Myc and Nanog show canonical activity in the undecided pan-ectodermal stem cells as well as in the neural domains, but the non-neural ectoderm has a negative activity score. The Bmp-signaling mediators Smad 1/5/9 are highly active in the non-neural ectoderm, whereas the Bmp/Tgfß inhibitor Smad 6 has high activity in the ventral midline of the developing neural tube. Tcf3, an activator of Wnt downstream signaling, which represses pluripotency in embryonic stem cells (203) showed high activity in the nonneural ectoderm. Also, Dlx5, Gata 2/3, known for their roles in the developing non-neural ectoderm are highly active. The cell cycle activators Id1/2/3 are active in the non-neural ectoderm and the undecided stem cells. The ventral neural tube also has high canonical activity of Foxa2, as well as Sox9 and FoxD3, and Tfap2A, which all play important roles in neural crest specification later on (2). The dorsal neural tube shows high downstream activity of Sall4, Zic1 and Hesx1, and Sox2, all known for their role in neural induction and stem cell maintenance (36, 44)(**Fig. 5A**).

**Figure 5.**
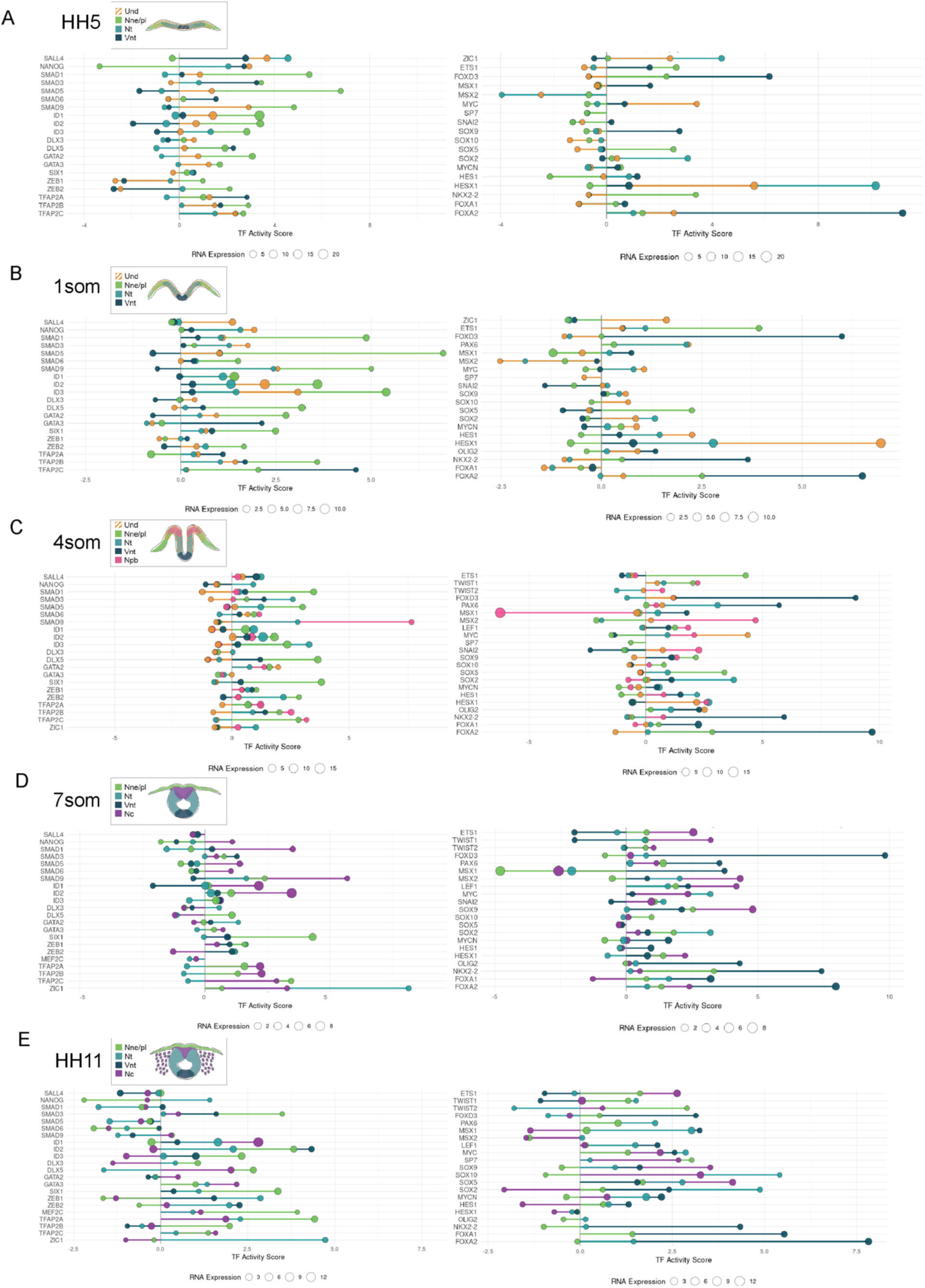
Canonical downstream activity scores both confirm relevance and bring novel insight of known lineage markers in the developing ectoderm. Lollipop plots show inferred activity scores and RNA expression levels for a curated set of genes that are known to be cell fate driving transcription factors (TFs). The figure only includes the TFs present in our regulatory resource conducted by the network analysis via decoupleR, thus excluding some known genes. Activity scores of known TFs at **A)** HH5, **B**) HH7 (1 som), **C**) HH8 (4 som), **D**) HH9 (7 som) and **E**) HH11. Bubble sizes correspond to RNA expression levels and are shown relative to the scale used within each individual plot Stem length reflects inferred activity.

#### Downstream activity of previously known genes at early neurula stage (1som)

The activity of the known genes in the early neurula is largely similar to the previous stage with some notable differences. Canonical activity of Zic1 is now solely in the undecided pan-ectodermal stem cells, which also show the highest activity of the Hesx1, which maintains stemness and self-renewal in the embryonic stem cells (36, 44) and the Tgfß-signalling mediator Smad3. The pre-placodal gene Six1 is now highly active in the non-neural ectoderm together with Ets1 and Sox5. Interestingly, Tfap2a is highly expressed in the non-neural ectoderm but shows a negative activity score whereas canonical activity is found in the other domains. On the other hand, Tfap2b and Tfap2c show canonical activity in all domains. The neural and the undecided stem cells have started to show activity of the neural lineage specifiers Pax6 (dorsal), and Olig2(ventral), whereas the Ascl1(aka Mash1), the master regulator of neurogenesis (204) shows high activity in the nonneural ectoderm at this stage. The neural plate border genes Msx1/2 activity patterns are interesting; canonical activity of Msx1 is only seen in the neural cell domains, whereas the undecided cells and the non-neural ectoderm present a negative activity score, and the Msx2 score is negative across all domains (**Fig. 5B**).

#### Downstream activity of previously known genes at mid-neurula stage (4som)

Again, the activity scores at the mid-neurula stage are largely similar to the previous early neurula stage. The early neural crest cells are rising at the neural plate border at this stage with notable canonical activity of Smad9 and the neural plate border marker Msx2. Also Twist 1/2, Sox9, Snai2, and TfAp2a/b/c, the non-neural ectoderm and early neural crest specifier genes that also prepare for EMT (2, 61, 83), as well as the Wnt-signaling mediator Lef1 show canonical downstream activity at the neural plate border and future epidermis. Notably, Msx1, which has the highest expression at the neural plate border shows a highly negative activity score, whereas canonical activity is present in the neural tube populations, suggesting a different role or repressed status. Interestingly, the CNS neural fate inducers Sox2 and MycN show canonical activity in the neural tube populations but, although expressed also in the future neural crest cells, the activity has a negative score suggesting a different role. The pro-neural genes Ascl1, En1 and Hes1 present high activity in the dorsal neural populations at this mid-neurula stage (**Fig. 5C**).

#### Downstream activity of previously known genes at the end of neurulation stage (7som)

At the end of neurulation, the activity patterns of the known lineage marker genes are similar to earlier stages. Nanog shows unique activity in the neural crest in line with previous studies on its role in neural crest pluripotency-like stemness (22, 205). The neural crest cells also have the highest activity of the Bmp-signaling mediator genes Smad1/5/6/9 as well as Lef1, and also Myc, which regulates the size and duration of the neural crest stem cell pool (95) and cell cycle genes ID1 and 2 also show high expression and canonical activity in the neural crest cells that are known to proliferate at this stage as they prepare for emigration EMT. The proneural genes Ascl1 and En1, which also plays a role in the development of the peripheral nervous system (206) were highly active in the neural crest cells. All three of the Ap2 genes and Sox9 show high canonical activity in the neural crest and in the non-neural ectoderm. Msx1 maintains its negative activity score in the rest of the ectoderm except the ventral neural tube that shows high canonical activity. Similarly, the canonical activity of Zeb2 (Sip1), which is shown to regulate adhesion during neural crest emigration and migration (107) is high in the neural tube populations but shows unique negative activity in the neural crest. High activity of Six1 is notable in the non-neural ectoderm, and Zic1 and Sox2 in the dorsal parts of the neural tube. The high activity scores of Foxd3, Olig, Nkx2-2 and FoxA1 and 2 are notable in the ventral neural tube together with the Wnt-mediator Tcf3 (**Fig. 5D**).

#### Downstream activity of previously known genes at post neurulation stage (HH11)

Finally, the gene activity patterns of the known genes remain similar as compared to earlier stages for the ventral neural tube and the non-neural ectoderm, although activity of Twist2 and Smad3 in the developing epidermis was new. On the other hand, neural crest cells, which are migratory at this stage, as well as the dorsal parts of the neural tube, which is reformed to establish the roof plate and overall integrity after neural crest cell emigration show changes in activity. Notably, Nanog and Myc show canonical activity in the dorsal parts of the neural tube, in line with our previous finding of a neural stem cell niche located next to the neural crest cells in the dorsal neural tube, which express pluripotency genes and may participate in the reconstruction after neural crest departure (22). Similarly, early neural lineage promoting genes Sox2, Zic1, MycN, Msx1, Hes1 and Zeb1/2, as well as En1 and Ascl1(36) showed high canonical activity. Bmp (Smads) and Wnt (Lef1) signaling mediators were no longer canonically active in the migratory neural crest cells, Id1 was highly expressed and active in these highly proliferative cells, and the SoxE (Sox9/10) SoxD (Sox5) transcription factors as well as Ets1, Myc,TfAp2a and Tfap2c showed high canonical activity, whereas TFap2b, Msx1/2 and the neural genes Sox2 and Hes1 showed notable negative activity scores, suggesting different roles of the neural genes in the developing CNS and the developing neural crest derived peripheral nervous system (**Fig. 5E**). Taken together, the analysis shows activity differences of known ectodermal markers between cell types as well as within a cell type at different developmental timepoints. While functional studies are required, these results bring valuable information on their putative roles in shaping lineage segregation during early development (**Fig. 5A-E**).

### Activity scores are not a readout of open chromatin status

Next, we asked whether the downstream activity score reflects the chromatin availability status of the enhancer binding region. In other words, do the negative scores correlate with closed chromatin and inability of the TF to bind? To test this, we used a previously published ATACseq data set from cranial HH9 chick embryo neural crest cells (207), which is equivalent to our HH9 neural crest sample, and analyzed chromatin binding availability of the 35 genes highlighted in the neural crest cells for either their positive or negative activity values at this stage (**Figs 6A-E**, list of individual ATACseq values of the 35 neural crest TFs from Figs 3C, 4D and 5D are found in **Supplemental Table 10**). The results show no difference between the occurrence of putative open binding regions for the TF motifs (**Fig 6A**), sum of total reads from all putative motifs that had greater than 99 reads per motif (**Fig 6B**), number of reads of the motif with the highest peak per each TF (**Fig 6C**), or the average of reads of the peaks of top 5 motifs per each TF (**Fig 6D**) between the TFs with negative vs positive downstream activity. Rather, both groups show similar gene to gene variability from low to high DNA binding activity on target motifs. Taken together, the neural crest example clearly demonstrates that the calculation of negative downstream activity cannot be explained by the inaccessibility of the TF to bind its enhancer but rather is an independent measure that reflects translational or posttranslational modifications that regulate their function, or alternatively, activation of a new non-canonical downstream cascade not recognized by the algorithm.

**Figure 6.**
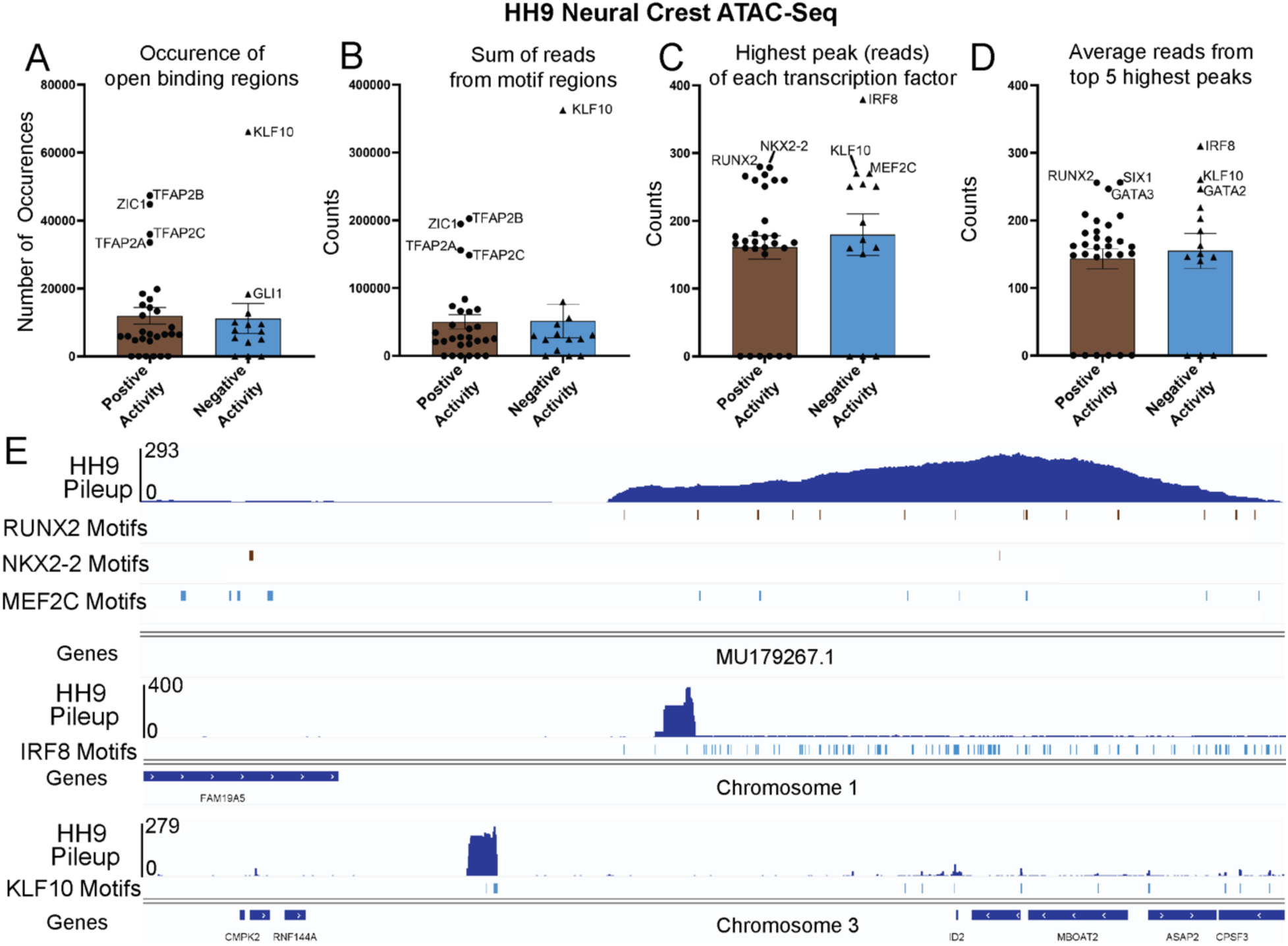
Quantification of ATACseq based chromatin binding on their putative motifs of the transcription factors that were highlighted for positive (brown) or negative (blue) gene activity in cranial neural crest cells at HH9 chick embryos. **A**) The sum of occurrences (open chromatin sites) for motifs of each transcription factor. **B**) The sum of reads from total motif occurrences (per gene) that had greater than 99 reads per motif. **C**) The highest peak count (number of reads) of each transcription factor (from a single motif) **D**). The average of reads of the five highest peaks (motifs) per each transcription factor. **E**) Examples of gene tracks with highest peaks identified at HH9 neural crest cells found in MU179267.1 (unannotated region in the chick genome), Chromosome 1, and chromosome 3. The figure shows where the motif occurrences are for the notable positive gene activity genes (brown bars) Runx2 and Nkx2-2, and the notable negative gene activity genes (blue bars) Klf10, Irf8 and Mef2c.

### Receptor–ligand analysis demonstrates the dynamic nature of interactions between tissues during head development

To explore how key developmental signaling pathways communicate between different tissues over developmental time, we next conducted a comprehensive analysis to map out the signaling interactions of the Wnt, Tgfβ, Bmp, Fgf, and Hh pathways between the ectoderm, endoderm and mesoderm (paraxial and lateral) at each embryonic stage (**Fig. 7A-E**). Utilizing CellChat (41, 208), we constructed an overview of cell–cell communication networks and visualized this using ligand–receptor interaction heatmaps. These heatmaps take into account the predicted strength of each cell type as a ligand source for the different signaling family members, and the strength of the probability of interaction with receiving tissues that express their receptors. It is worth noting that the analysis reveals interactions that are most probable to occur based on gene expression, and some genes that are lowly expressed, yet biologically relevant, may be filtered out from the results (**Fig. 7**). First, the results on WNT signaling (**Fig. 7A**) highlight a few interesting patterns; the earliest probable interactions occur at the 1 somite stage, when signals emanate broadly across all cells bar the endoderm and notochord. As development progresses, a shift occurs with the ligands being predominantly expressed by the neural crest and the neural tube but, interestingly, primarily targeting the mesoderm. At mid-neurula stage, Wnt signaling is still received abundantly also by the neural ectoderm, but the notably high scale due to strong mesodermal response fails to detect additional potentially more subtle responses in the ectoderm at the later stages. FGF signaling, on the other hand, is characterized by its clear dominance in signaling between ectodermal cell types during the first three stages. By neural tube closure, while nearly all cells actively participate in both sending and receiving signals, the ventral neural tube remains as the highest predicted source and receiver of FGF signaling (**Fig. 7D**). Furthermore, Bmp signaling follows a distinctive trajectory in terms of its source tissue; initially (HH5/1 somite), signals are predominantly issued from the endoderm, notochord and, to a lower extent, from the non-neural ectoderm and likely received by all cell types. Signaling is transcribed predominantly by the notochord with a gradual shift to the mesoderm as the principal signal source over time for predominantly ectodermal cells (**Fig. 7C**). In contrast, TGFβ signaling showcases a consistent pattern, with the highest signal originating from the notochord and directed towards all cells across every embryonic stage (**Fig. 7B**). Similarly, the notochord and the ventral neural tube serve as consistent signal sources of the HH signaling pathway across all stages, however, the recipient profile evolves from all cells, to just the neural tissue and paraxial mesoderm at later stages (**Fig. 7E**).

**Figure 7.**
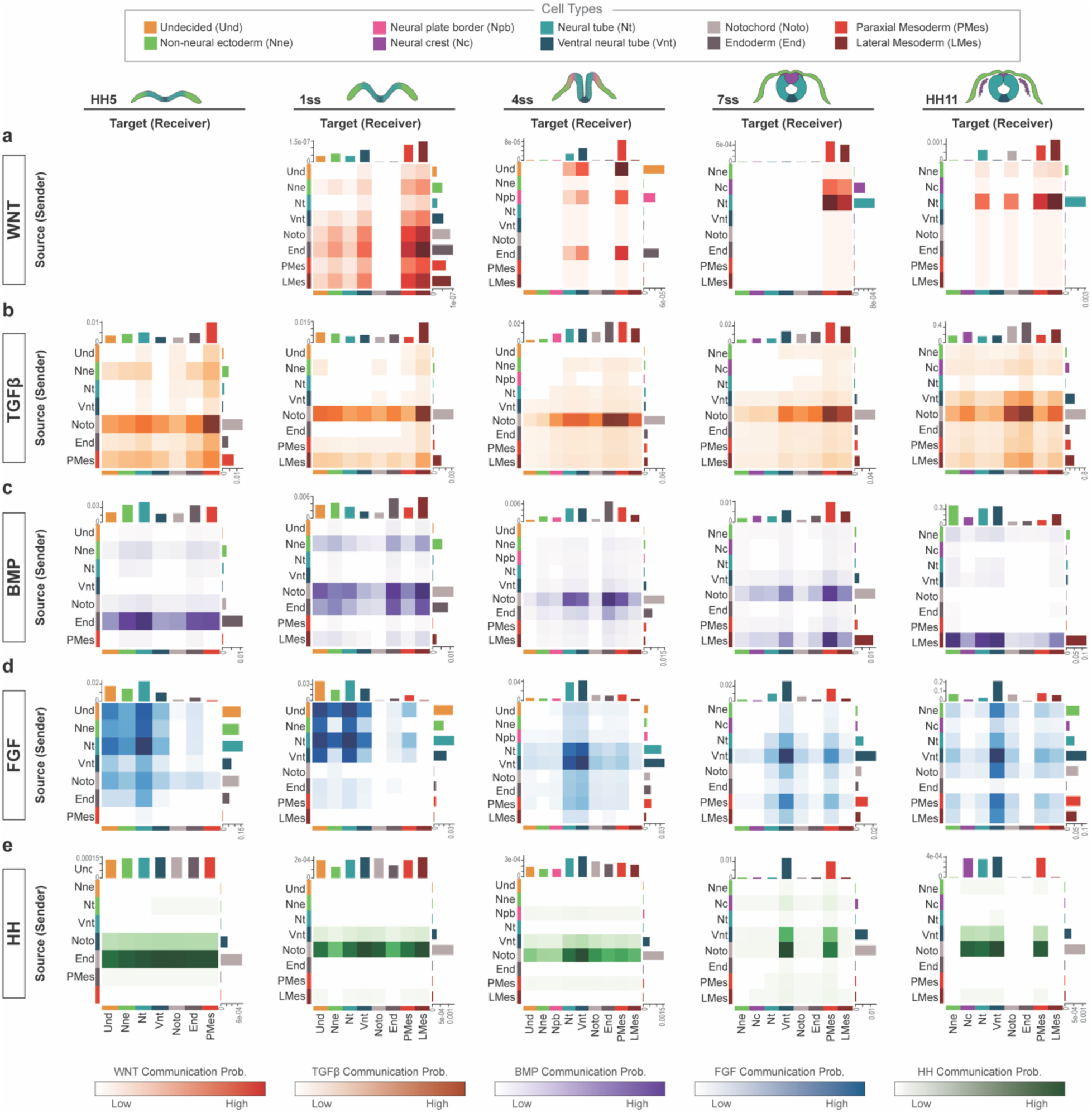
Cell-to-cell signaling prediction (Cell Chat) of the developing head confirm established mechanisms and reveal novel signaling sources and interaction details between and within germ layers. **A)** At early neurulation stages, Wnt signaling is produced by all germ layers and particularly by the endoderm and notochord and mainly received by the mesoderm as well all ectodermal cell types. At mid-neurula stage, the endoderm as well as the undecided stem cells and the neural plate border produce Wnt, which is further restricted to the neural crest and the dorsal neural tube at the end of and post neurulation stage and mainly received by the mesoderm. **B**) Thoughout ectodermal patterning, Tgfb signal is secreted mainly by the notochord and paraxial mesoderm and received mainly by the mesoderm and to a lesser extent by all germ layers during early stages and by endoderm, notochord and the neural tube and neural plate border at mid and end of neurulation stages. **C**) The strongest source of BMP signaling after gastrulation is the endoderm, which is received by itself, the undecided stem cells, dorsal neural plate, the non-neural ectoderm and mesoderm. In the early neurula (1 som) the non-neural ectoderm and notochord also produce signals, and notochord together with the lateral mesoderm become the strongest source at late neurulation stages, which is received by all germ layers, particularly the ventral neural tube and the paraxial mesoderm. After neurulation, the main BMP signaling source is the paraxial mesoderm with lower input from the nonneural ectoderm, and the signals are mainly received by the ectoderm. **D**) The strongest interactions of FGF signals during ectoderm patterning are within the ectoderm, which switches from interactions between all the cell types at early neurulation to the neural tube populations at mid-neurula stage to mainly the ventral neural tube signaling to itself at post-neurulation stage. **E**) Highest Hedgehog signaling is initiated form the endoderm after gastrulation and switches to the notochord and ventral neural tube during rest of the neurulation, and the neural plate border and crest also send signal at premigratory stages. All cell types receive signal until mid-neurulation, which shifts to ventral neural tube and paraxial mesoderm at the end of neurulation, as well as to the dorsal neural tube and the neural crest post neurulation. (The specific ligand -receptor pairs are shown in **Supplemental figures 3-5**).

The full ligand–receptor analysis reveals the individual molecule pairs with the strongest predicted interactions, providing a myriad of novel information, details of which are presented as dotplots (**Supplemental Fig. S3–5).** There has been a wealth of work produced on signaling pathways by the chick research community, and as such, we validated the vast majority of these predicted interactions by verifying the correct spatiotemporal expression from *in situs* from published literature, details of which are included in **Supplemental Figure S6** (209–229), which also greatly increase confidence of the novel interactions found in our data set. In certain instances, we felt that there was not the data to support our findings, and so we conducted HCR ISH (**Supplemental Fig. S7**) to validate these predictions. For example, *Fgf6* has been previously reported as not expressed before HH14 (224), whereas our predicted interactions identify *Fgf6* as likely signaling at HH7 from undecided ectodermal cells, the non-neural ectoderm and the neural tube, and this signaling from neural tissue is likely to persist through to HH9, also accompanied by signaling from the notochord, paraxial mesoderm and endoderm with some likelihood (**Supplemental Fig. S5**). As such, using HCR ISH we have confirmed that this gene is expressed in the ectodermal tissues at HH7 and HH9 as predicted, with some expression in the mesoderm and endoderm at HH9 (**Supplemental Fig. S7H–I**). In sum, we were able to validate all the strong interactions predicted by the ligand-receptor analysis. However, our approach is not able to address the biological relevance of previously reported interactions which may have not been picked by the analysis based on their transcriptional profile. A summary of this analysis is depicted in the schematic in **Fig. 8**. In sum, our results offer fresh insights into signaling strengths of various ligands between different tissues during embryogenesis over time.

**Figure 8.**
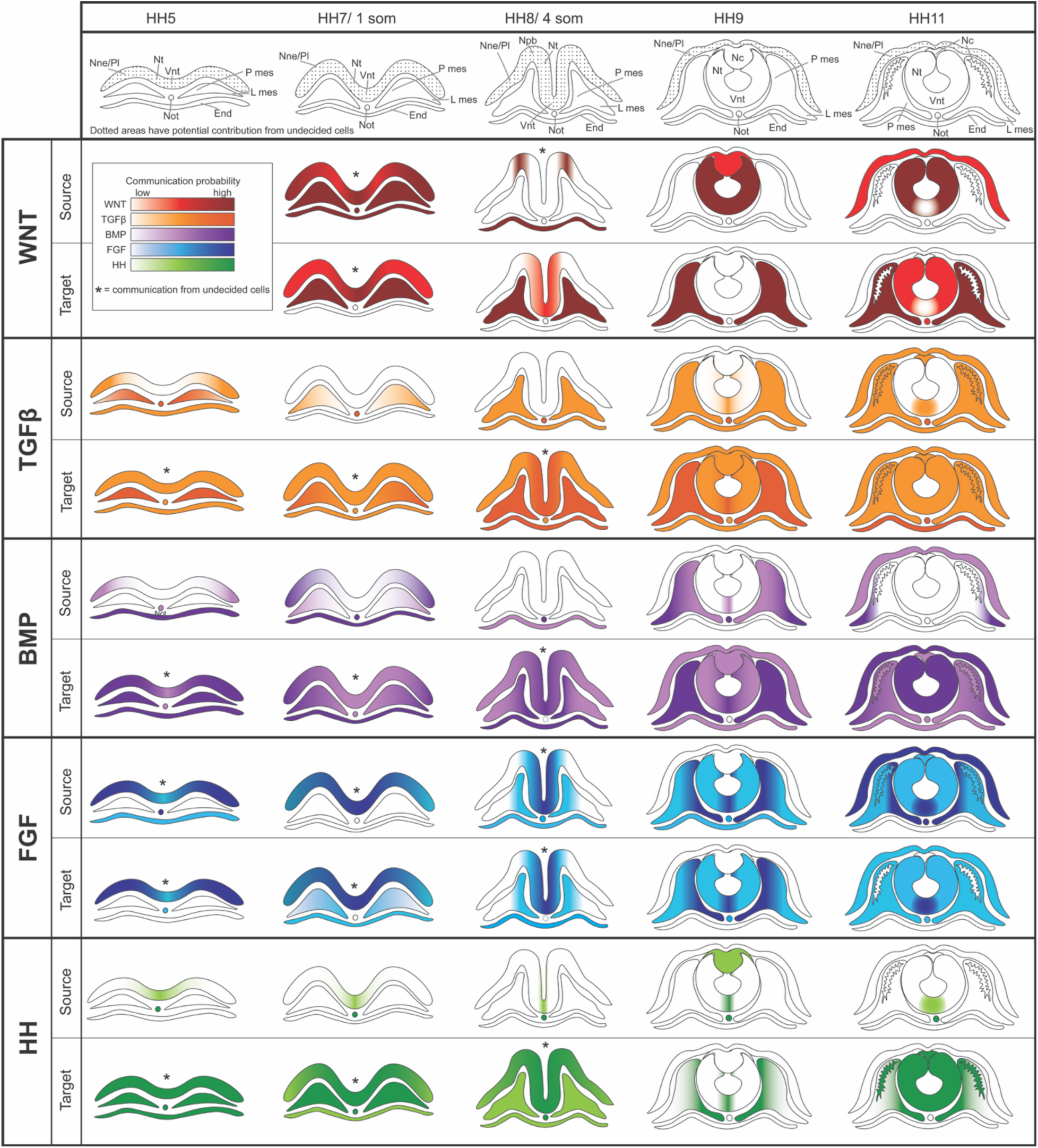
A schematic of the main signaling pathway interactions in the neurulating embryo at midbrain level.

## Discussion

Embryogenesis is orchestrated by the coordinated actions of activating and inhibitory cues from a limited set of conserved signaling pathways (Levine and Brivanlou, 2007; Rogers and Schier, 2011; Suzuki and Suzuki, 2020; Schauer and Heisenberg, 2021). These early signaling events trigger distinct transcriptional programs across diverse cell populations and states, resulting in the expression of thousands of genes. Despite over a decade of large-scale bioinformatic screening, our understanding of cell type–specific fate determination remains centered on a relatively small group of transcription factors identified decades ago. A key limitation of such analyses is that novel candidates are often overlooked because many genes appear ubiquitously expressed, obscuring subtle but functionally relevant differences. This challenge has particularly restricted insights into the cell type–specific roles of lowly expressed epigenetic and post-translational regulators. Moreover, while transcription factors have been extensively studied within individual cell types, their expression in neighboring domains—and potential context-dependent functions—remain poorly understood. Although scRNAseq has become increasingly widespread rapidly increasing the amount of publicly available data sets, the field now faces the challenge of developing more sophisticated analytical frameworks to extract deeper biological meaning, as RNA abundance alone provides an incomplete measure of protein activity and regulatory state.

To circumvent these challenges, we employed **TF Activity Inference** (28), a recently developed computational framework designed to estimate TF activity from the expression patterns of canonical target genes rather than relying exclusively on transcript abundance, thereby providing a more functionally informed representation of regulatory dynamics derived from scRNA-seq data. The results highlighted important patterns. First, the canonical downstream activity of transcription factors and epigenetic modifiers does not correlate with RNA expression levels. Our results defined several genes, including known lineage drivers like Tfap2a of the non-neural ectoderm, which displayed differential canonical activity despite similar RNA expression levels in both neighboring tissues as well as in the same lineage across different developmental timepoints, suggesting at least context-dependent nuances in their regulatory input but potentially also separate roles. The genes with highest activity consisted of both known lineage and pluripotency and proliferation driving factors like Sall3, Hex1, Foxa2, Hand2, Zic1 and Id3 as well as epigenetic modifiers like Ssrp1, and Ncor2 (**Fig 2**). Together, our results highlighted thirty epigenetic modifiers, seven RNA binding proteins, and fourteen cell cycle regulators not known to play cell type or developmental stage specific roles during ectoderm patterning before. Futhermore, cross referencing of the results with ATACseq based TF motif availability and occupancy revealed that the downstream activity algorithm does not correlate with TF binding availability to its enhancer DNA region, as TFs with negative activity showed no difference in their variance of open chromatin binding sites as compared to TFs with positive activity scores (**Fig 6**). Thus, the TF Activity Inference algorithm presents an additional, independent and credible tool to evaluate transcription factor activity from scRNAseq data, as predicted canonical activity of known ectodermal fate driving TFs was largely in line with current literature.

Second, out of the hundreds of expressed genes, we identified a handful of genes that were consistently active in their respective cell populations during all stages of ectoderm patterning and at post-neurulation building a framework of essential lineage guiding activity (**Fig 3**). In line with existing knowledge from several decades, all cell types included a few known transcription factors of the respective lineage (Sall4, Sox2 for undecided stem cells; Dlx5, Six1 for non-neural ectoderm; Tfap2a, Twist1 and Sox10 for NC; Sox2, Pax6 for dorsal and Foxa2 for ventral neural tube). However, most of the highlighted canonically active genes were novel in this context. Several epigenetic modifiers were included, of which some were shared between cell types like Brca1, which showed canonical downstream activity in stem cells and the dorsal neural tube, Ncor2 which was shared by dorsal neural tube and the neural crest, and Ssrp1 that showed activity in the neural crest and the ventral neural tube, offering a valuable resource for understanding how chromatin is regulated during distinct ectodermal cell fate transitions. Future studies will be important for elucidating the interplay among these factors.

Notably, continuous canonical activity of Beta-catenin (Ctnnb1), which regulates cell adhesion, cytoskeletal dynamics and migration, as part of but also independent of Wnt signaling (84), was highlighted in all cell types except for the stem cells, suggesting that it plays a pivotal role in ectodermal morphogenesis and formation of tissue barriers. Interestingly, hypoxia and glycolysis related regulators were highlighted for their consistent canonical activity in several cell types: Hif1a was detected in both the non-neural ectoderm as well as the ventral neural tube, which also showed high activity of Ybx1, a multifunctional DNA and RNA binding protein, which directly binds to Hif1a mRNA to stabilize it under hypoxia in hepatocellular cancer cells (115, 182). Furthermore, Enolase 1, which is a glycolysis metabolite that can be induced by hypoxia, of which an alternative isoform can also act as a transcription factor (111) showed high activity and expression throughout dorsal neural tube development. Whether and how hypoxia is involved in governing ectodermal patterning will be an interesting future direction.

The results also highlighted unprecedented details of signaling pathway mediators. The Notch signaling receptor Notch1, that inhibits expression of proneural genes and maintains neuroblasts in an undifferentiated state (230) showed negative values in the pan-ectodermal stem cells. Furthermore, the Shh activator Gli1 presented negative activity in the nonneural ectoderm, and both Bmp signaling downstream activators (Smad5/9) and Gli3, a Shh signaling modulator (activator in the presence and inhibitor in the absence of the signal)(119) showed negative values in the ventral neural tube, in line with the opposing roles of ventral Shh and dorsal Bmp inductive signals during neural tube patterning. Finally, the genes highlighted for their continuous negative activity through ectodermal patterning in each cell type mainly consisted of broadly expressed epigenetic and cell cycle regulators, and the analysis thus brings insight of their cell type specific differences.

Third, many cell cycle regulators and epigenetic modifiers are ubiquitously expressed, which has prevented them from being highlighted in conventional gene expression-based analyses. To distinguish between potential developmental stage or lineage specific differences in their functional activity we applied downstream activity inference to identify genes at each developmental stage that presented either exclusive activity (beyond 0) in only one of the cell types, or alternatively, presented a gradient of activity (beyond 1) in only one cell type and lower in others (**Fig. 4**). Many of the highlighted genes from the analysis were epigenetic modifiers and cell cycle regulation and DNA repair related genes that have not been studied in this context before, bringing valuable new information about the chromatin landscape changes during ectodermal lineage segregation and cell fate commitment. Notably, the results revealed unique activity of Prdm1 and Smarca5 in the non-neural ectoderm, Ncor2, Brca1 and Dnmt1 in the dorsal neural tube, Kat7 in the ventral neural tube, and Dnmt3B, Smarcc1, Znf143, Mta1 and Ncoa1 in the NC cells during the course of ectodermal patterning.

Furthermore, some genes presented significant differences between activity scores in different cell types of which the activity of Prdm1 between the non-neural activity and the negative score of the ventral neural tube was most notable. Furthermore, apart from some known lineage promoting transcription factors, some downstream effectors of signaling pathways were highlighted, which brings detailed insight about the specifics of downstream activity mechanisms. At the earliest stage HH5, these included the activity of the non-canonical Wnt downstream mediator Tcf12 and the Yap signaling mediator Tead in the non-neural ectoderm as well as the sonic Hedgehog signaling inhibitor Gli3 (108) activity in the developing dorsal parts of the neural plate. In the early neurula, Notch1 and the Bmp inhibitor Smad6 (142) were highlighted in the non-neural ectoderm. Finally, at the end of neurulation, activity of the Bmp activator Smad5, which was also continuously active in the pan-ectodermal stem cell population from post gastrulation to mid neurulation, was unique to the neural crest cells, which additionally showed unique activity of the retinoic acid receptor Rarb. Furthermore, the Notch signaling downstream activator and proneural gene Hes1, Wnt-signaling mediator Tcf3 and TgfB-signaling mediator Smad3 and Irx1, which is linked to dorsoventral patterning of the central nervous system (199), presented unique canonical activity in the ventral neural tube.

Finally, we used downstream activity inference to study the potential differences in the activity of known genes that have been indicated in published gene regulatory networks for specific lineage driving roles (2, 36, 135, 231). In conclusion, the majority of genes previously linked to specific cell types displayed canonical activity in those cells at some developmental stage, supporting the robustness of our analysis. Of note, as TF activity scores can only be calculated for genes that are present in the input list that is supplied to the network analysis, not all TFs could be assigned activity scores. Furthermore, since the TF-target interactions in CollecTRI are derived from human data, performing activity inference across species further limits which TFs have compatible target information. For these reasons, some widely used markers like Pax7/3, Sox8, Sox1/3 are left out. Noteworthy exceptions in canonical activity included the neural plate border genes Msx1 and Msx2, which have also been suggested to play compensatory roles with each other. Msx genes are highly expressed in the neural plate border and the premigratory neural crest cells and knockdown studies have shown they play essential functional roles in neural crest development as a mediator of Wnt and Fgf signaling and pro-apoptotic activity largely via co-binding activity to other transcription factors rather than direct enhancer binding (61, 92, 135, 232–235). Our results suggest a different downstream role for Msx1 in the neural tube and particularly its ventral portion, which shows positive canonical activity scores, as compared to highly negative activity in the neural crest. On the other hand, Msx2, which is also abundantly expressed in the neural plate border region already at stage 5 and later in neural crest cells, showed negative activity in all cell types at early neurula stage, but switched to canonical activity in the neural crest at mid-neurula stage, which switched back to negative values in the migratory neural crest - thus reflecting different time dependent roles during neural crest development. Taken together, the MSX1/2 example suggests an alternative, cell type and developmental stage specific functional mechanism for these transcription factors that cannot be explained by enhancer binding availability differences. Similarly, MycN, a key driver of CNS neural stem cells that is amplified in the neural crest derived pediatric cancer neuroblastoma (236, 237), showed interesting differential activity patterns: while the results show canonical activity in the ventral neural tube, the activity scores were negative in the developing neural crest indicating different roles, which may be of significance for therapeutic planning of neuroblastoma. Similarly, the canonical activity of Sox2 was only presented in the developing neural tube but not in the neural crest cells, and the activity scores of the pro-neural gene En1 (120, 206) shifted from high canonical activity in the premigratory neural crest cells to negative activity in the migratory stage, where high canonical activity was shown in the dorsal neural tube instead. The high canonical activity of Smad9 stood out in the premigratory neural crest /neural plate border cells warranting further functional investigation of cell type specifics of downstream activity of Bmp signaling in the ectoderm. Finally, the known neural crest specifier gene FoxD3 presented high canonical activity in the ventral neural tube and, in comparison, the neural crest downstream activity was modest. Furthermore, the different nuances of ID-gene activities between cell types and developmental stages reflects specifics in cell cycle activation during ectoderm patterning. In sum, the results underscore the context-dependent deployment of downstream activators and delineate which cell types and developmental stages exhibit shared versus distinct patterns of downstream gene activity opening multiple interesting future research avenues.

One of the debated questions of neural crest development is the origin of the pluripotency-like characteristics of the neural crest cells – whether they stem from the common stem cell population shared with the rest of the ectoderm (22), or a transcriptionally distinguished neural crest stem cell lineage separate from the rest of the ectoderm (238), or as a *de novo* emerging population of the neural plate border region (46). Our results from the RNA velocity prediction indicates that the neural crest cells stem from the same population of undecided pan-ectodermal stem cells as the rest of the ectodermal domains, which transition into the neural plate border cells before final specification to premigratory neural crest cells. Furthermore, the canonical activity of the pluripotency genes Nanog and Sall4 throughout the early stages of ectoderm development after gastrulation is in line with our previous work where we found that until mid-neurulation stage, the entire ectoderm contains pluripotent-like stem cells, which are transcriptionally undecided between future ectodermal fates (22).

Our results also provide detailed insight on signaling pathways through a growth factor–receptor signaling atlas that quantifies the activity strength of each pathway based on ligand expression levels in source and receiver cell types throughout the stages of early development at the midbrain axial level (**Fig. 7-8, Supplemental Fig. S3–6**). Previous understanding of signaling pathways has been heavily informed by experiments involving the addition of ectopic growth factors (239, 240), which often do not employ the correct *in vivo* ligands or elucidate the endogenous sources or specific temporal expression patterns of these factors. Consequently, the developmental biology field frequently relies upon a generalized understanding of signaling activity, such as the differential roles of Bmp, Fgf and Wnt signaling in ectoderm and neural plate development, without precise spatiotemporal expression knowledge of the signaling molecules and their downstream mediators of the respective families. With the unique opportunity to measure signaling pathway activity simultaneously across all cell types, our work aims to bridge this gap, offering a more nuanced, detailed insight into the specific temporal and spatial expression patterns of specific subtypes of signaling molecules within various families.

The Cell-Chat results thus allowed us to follow overarching, large signaling entities relevant to embryogenesis, which revealed the most dominant sources of respective signaling pathways and provided interesting novel details not acknowledged before. For instance, at HH5, Bmp5 signal produced by the endoderm emerges as the most likely significant Bmp signaling source to the non-neural ectoderm (**Supplemental Fig. S4**). This is in contrast to the current view where ectodermally expressed Bmp4, which is often referred to as the sole inducer of the non-neural ectoderm fate (241), but which according to the prediction, has a much lesser impact. It is important to acknowledge that our receptor–ligand analysis employs prediction on normalized and relative expression levels and thus do not *per se* contest any shown interactions within, which may be left out from our results due to the strict stringency of our analysis. Despite these constraints, we were able to verify all the strong interactions predicted by the analysis across germ layers by *in situ* hybridization (both here and previously published; Supplementary **Figs. S6****-9**), which greatly increases the credibility of the previously unknown, interesting interactions that are worth further exploring to understand the whole picture of signaling events in the developing head. Multiomics studies will be essential for future discoveries to understand the complexity of co-regulatory mechanisms that guide early development. In sum, even though future functional experiments are required to validate the accuracy of the individual interactions, the consistency of the matching results provided by the ligand–receptor probability analysis together with the gene expression and downstream activity analyzes, such as matching downstream activity results of Smad-proteins downstream of Bmp and Tgfb signaling as well as by knowledge from previous studies (1, 11, 14, 45, 240, 242–245) together provide a convincing source for the understanding of unstudied interactions. To our knowledge, studies that gather comprehensive information of an entire developing organism on a specific axial level have not been performed, and our high-resolution analysis thus provides a new level of molecular accuracy regarding activation of signaling molecules and their putative receptor pairs as well as novel insights into the main source tissues of signaling molecules in the developing head.

## Material and methods

### Chicken Embryos

Fertilized chicken eggs were sourced from two distinct locations: the University of Connecticut (UCON) poultry farm (CT, USA) and Lassilan Tila (Tuusula, Finland). The eggs were incubated at 37°C and closely monitored to attain the desired developmental stage, following the Hamburger and Hamilton (HH) classification system, as illustrated in Fig. 1a for precise stage definitions. All embryos utilized in this study were younger than three days of age, which aligns with regulations stipulated in The Finnish Act on Animal Experimentation (62/2006) and guidelines outlined by The Institutional Animal Care and Use Committee (IACUC) of NIH. Ethical approval for the use of such embryos was not required.

### 10X Chromium scRNA-seq

For single cell sequencing, cDNA from chick embryo samples (HH5, 1som, 4som and 7som) was generated and sequencing libraries were prepared as previously described (Pajanoja et al., 2023). Additionally, this study introduces a novel dataset of HH11 stage chick embryos (ranging from 13–14 somite stage). Briefly, midbrain slices covering all germ layers were dissected from the five developmental stages using micro scissors. These slices were collected from three to six embryos per sample (with two replicates for each stage except for only one sample for HH11). The tissue was then dissociated into a single-cell suspension using the Miltenyi Biotec multi-tissue dissociation kit (kit 3,130-110-204, Miltenyi Biotec).

Sequencing was carried out using the 10x Genomics Chromium Single Cell 3’RNAseq platform at the FIMM Single-Cell Analytics unit, with support from HiLIFE and Biocenter FinlandLibrary preparation and sequencing were performed using the Chromium Next GEM Single Cell 3’ Gene Expression version 3.1 Dual Index chemistry on an Illumina NovaSeq 6000 system. The sequencing run included read lengths of 28bp (Read 1), 10bp (i7 Index), 10bp (i5 Index), and 90bp (Read 2). All eight samples were sequenced together in a single run. Fastq files were generated using the 10x Cell Ranger (v7.0.1) count pipeline for alignment, filtering, and UMI counting. Finally, the Fastq files were mapped to the chicken genome using a custom GRCg6a Gallus gallus genome annotation file. The sequencing of the HH11 stage samples was conducted at the NIDCD/NIDCR Genomics and Computational Biology Core (on Illumina NextSeq 2000 system; read lengths of 26bp (Read 1), 8bp (i7 Index), and 93bp (Read 2)), adhering to the same sequencing standards to ensure uniformity and comparability across all datasets.

### scRNA-seq Analysis

In alignment with our previous analysis (Pajanoja et al., 2023), data cleaning, normalization, and scaling procedures were consistently applied to the HH11 stage chick embryo sample. Specifically, we employed the Seurat package (v3.0.1) in R for data analysis. The SoupX and DoubletFinder packages were employed to filter out ambient RNA and doublets, respectively (**Supplemental Fig S1A**). Subsequently, we evaluated the presence of batch effects by comparing the gene expression profiles between HH11 stage chick embryo sample and the previously sequenced samples. Remarkably, our analyses, which included various visualization techniques such as PCA plots, t-SNE plots, and hierarchical clustering, revealed no discernible batch-related discrepancies or systematic differences in gene expression across the entire dataset, and no additional integration package was required (**Supplemental Fig. S1B, C**).

The analysis initially started by addressing each of the five developmental stages individually. The standard Seurat workflow was applied for scRNA-seq data analysis, with careful consideration given to data quality and consistency. Exclusion criteria were applied to cells, resulting in the exclusion of those in which an extreme number of genes were expressed, exceding either 4000 or 5500 genes, and those with a mitochondrial content surpassing 0.4%. Following this initial data filtration step, the gene expression values were normalized and scaled using the ‘NormalizeData’ and ‘ScaleData’ functions, respectively. Dimensionality reduction via principal component analysis (PCA) was performed using the ‘ElbowPlot’ function, with the retention of the top 20 principal components (dims 1:20), while employing a resolution of 0.6. Clustering of cells was generated and visualized through UMAP plots. For the purposes of cluster annotation enhancement, manual annotations were based on differentially expressed genes within each cluster. These genes were identified utilizing the ‘FindAllMarkers’ function, with log-fold change thresholds of 0.2 and minimum percent thresholds of 0.25. Furthermore, annotation of each sample was carried out in accordance with the germ layers, which encompassed the ectoderm, endoderm, and mesoderm, and a discrete population of cells corresponding to the notochord (**Fig. 1B**, **Supplemental Fig. S1–2**).

### Ectodermal Cell Populations

’Ectoderm’ clusters for each of the five stages were subsetted and individually analyzed (**Fig. 1A**). Following the cell type annotation process, the results from all developmental stages were merged and re-clustered. Subsequently, differential expression analysis was applied to uncover variations not only among cell types but also across developmental stages within the ‘Ectoderm’ clusters. This analysis was conducted using the ‘FindAllMarkers’ function, with default settings for the Wilcoxon rank-sum test, log-fold change threshold set at 0.2, and a minimum percent threshold of 0.25. Additionally, cell cycle phase percentages were computed using built-in functions provided by Seurat, further enhancing our understanding of cellular dynamics within the dataset.

To enable in-depth characterization of cellular populations within each developmental stage, Seurat’s ‘ident’ feature was utilized to assign distinct labels to cells in the merged dataset (**Supplemental Fig. S2A**). These labels were systematically combined with stage information and cell type designations, resulting in the creation of 21 unique identifiers for each specific stage and cell type it represented. As a result, this provided the means to systematically compare and contrast molecular signatures and expression profiles across cell types and stages, allowing for a thorough side-by-side analysis of the entire dataset.

### Transcription Factor Analysis, Pseudotime and RNA Velocity

Comprehensive lists of human and chicken transcription factors (TFs) were accessed from the KEGG BRITE classification system (https://www.genome.jp/kegg/brite.html). Human gene lists were then converted to chicken genes, resulting in the identification of a total of 903 TFs within our dataset. The incorporation of these TFs into our dimensionality reduction and clustering workflow allowed for the establishment of connections between clusters. DE genes were then calculated using this Seurat object exclusively containing TFs (Logfc.threshold= 0.2, min.pct= 0.25).

Next, all genes were re-integrated into the Seurat object while preserving the newly acquired UMAP embedding and was subjected to pseudotime analysis using Monocle3. First, HH5 was set as the starting stage, then the proximity of these cells to the principal graph’s vertices in UMAP space was calculated to select the root node for the pseudotime trajectory. Trajectory analysis was carried out using ‘learn_graph’ function. Cells were systematically ordered in pseudotime, with reference to the earliest principal node calculated during the analysis.

RNA velocity analysis was conducted using the same UMAP embedding. Loom files for each five developmental stages were created using the Velocyto.py command line tool. Ratios of unspliced to spliced RNA were computed utilizing the scVelo dynamical modeling pipeline and RNA velocity embedding was drawn on the UMAP based on the analysis.

### Transcription Factor Activity Analysis

Transcription factor activity scores were computed using decoupleR with the CollecTRI human regulatory resource. Log fold change values from differentially expressed genes across 21 unique idents (**Supplemental Fig. S2A**) were used as input. CollecTRI gene–TF relationships were applied to compute TF activity scores following the decoupleR framework. For downstream analyses, TFs were retained only if their corresponding RNA expression values were more than zero.

For gene ontology enrichment, the top 30 highest and lowest TF-activity genes werw selected separately and analyzed using DAVID (Biological Process). Significance was assessed with Fisher Exact test and adjusted for multiple comparisons using the Benjamini–Hochberg method.

### ATAC- Seq Analysis

The raw ATAC-sequencing data of HH9 cranial chicken neural crest was previously published (207) and FASTQ files were downloaded through the Gene Expression Omnibus: GSE126880. Reads were aligned on the ENSEMBL chicken genome (GRCg7b) using STAR (246). Peak calling was then performed using MACS version 3.0.3 in with -f BAMPE, and -q 0.01. Transcription factor motif occurrences were analyzed on the open chromatin regions using FIMO from MEME Suite version 5.5.9 (https://doi.org/10.1093/nar/gkv416; https://meme-suite.org/meme/tools/fimo). Motif PFMs were downloaded from the 11^th^ release of JASPAR (https://jaspar.elixir.no/downloads/). GFF files of each transcription factor of interest was downloaded from FIMO, which was used with bedtools intersect (with a cutoff of 5bp or higher overlap) to generate a table of reads for each motif occurrence. Gene tracks were visualized using Integrative Genomics Viewer (https://www.nature.com/articles/nbt.1754; https://igv.org/doc/desktop/).

### Receptor–Ligand Analysis

For the receptor-ligand analysis, all cell types from all germ layers were considered, including the mesoderm cell clusters in each stage which were annotated as ‘paraxial’ and ‘lateral mesoderm’, and each developmental stage was analyzed individually. The analysis was conducted using the CellChat package and the highest recommended stringency value, which leverages existing human ligand–receptor interaction database in the absence of an available dataset based on chicken protein–protein interactions. This decision was informed by the high degree of evolutionary conservation observed in key signaling molecules and pathways across vertebrates, coupled with the comprehensive coverage of human ligand–receptor pairs in the database. It is important to note that while this method offers valuable insights, the interpretations were made with careful consideration of the potential limitations and the evolutionary context between the species. Outcomes of the analysis were presented, highlighting the identified ligand-receptor pairs and key signaling pathways involved in the communication network using the “netVisual_heatmap” function.

### Whole mount Fluorescent *In Situ* with Hybridization Chain Reaction

Chicken embryos were fixed in 4% paraformaldehyde, washed with PBS/0.2% Tween (PBST), dehydrated in methanol, and stored at -20 °C. HCR split-initiator probe sets were purchased from Molecular Instruments (www.molecularinstruments.com) for chicken *TPI1* (NM_205451.1), *FOXK2* (NM_001395217.2), *ACTR3* (NM_204307.2), *COBL* (XM_040695853.2)*, BMP4* (NM_205237.4)*, WNT3A* (NM_001398207.1)*, WNT4* (NM_204783.2)*, PTC1* (NM_204960.3)*, FGF4* (NM_001031546.3)*, FGF6* (XM_001232070.5). In situ HCR v3.0 with split-initiator probes was done according to whole-mount chicken protocol (247) with hybridization overnight at 37°C and amplification overnight at room temperature. Following amplification, embryos were washed with 5xSSCT, stained with DAPI, and post-fixed with 4% PFA for 15 minutes at room temperature. Embryos were embedded in gelatin, sectioned at 20 μm and imaged using an Andor Dragonfly spinning disk confocal microscope.

## Data Availability

The scRNA-seq data for the chick cranial developmental stages is deposited in the NCBI GEO database under accession number GSE221188 and the neural crest ATACseq data set (207) is available at GSE126880. Analysis code used in this study is available upon request. Please contact for access to the code.

## Author Contributions

CP performed all scRNAseq based analysis. JR performed all Cell Chat validations. ET performed ATACseq analysis. CP, JR, ET and LK prepared figures. LK wrote the paper with input from JR, CP and ET.

## Acknowledgements

We thank Daniel Martin for helpful discussions on the methodology of the manuscript.

---

**Supplemental Figure 1.**
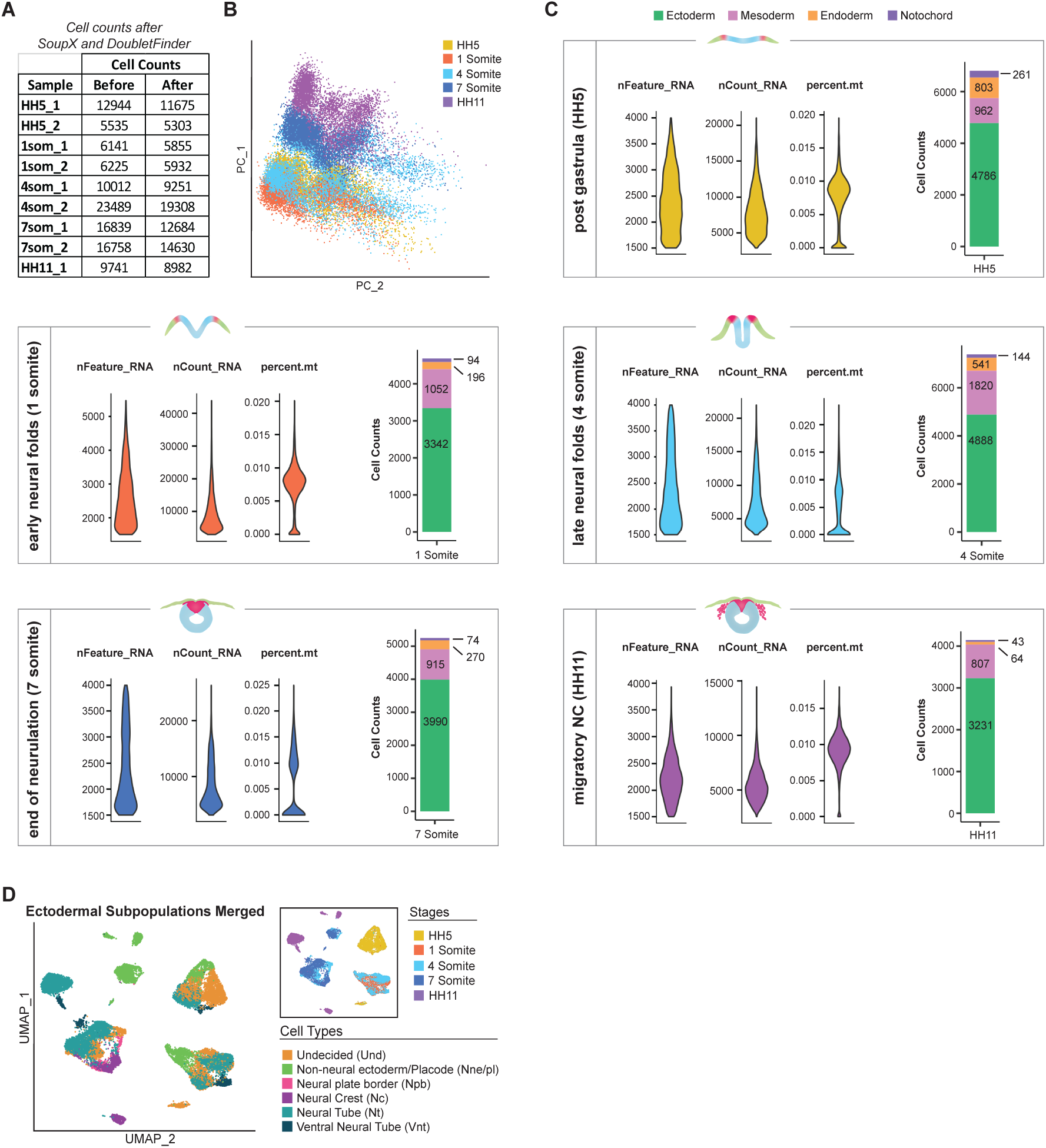
**A)** Cell counts of the scRNA-seq samples pre- and post-cleanup using SoupX for ambient RNA removal and DoubletFinder for doublet identification. **B)**, PCA analysis of merged samples, demonstrating no apparent batch effects across the dataset. **C)**, Violin plots for each of the five developmental stages, illustrating nFeature_RNA, nCount_RNA, mitochondrial content, and cell distribution across germ layers. **D)** Merged ectodermal subsets from all five stages, color-coded by cell type and stages separately.

**Supplemental Fig 2.**
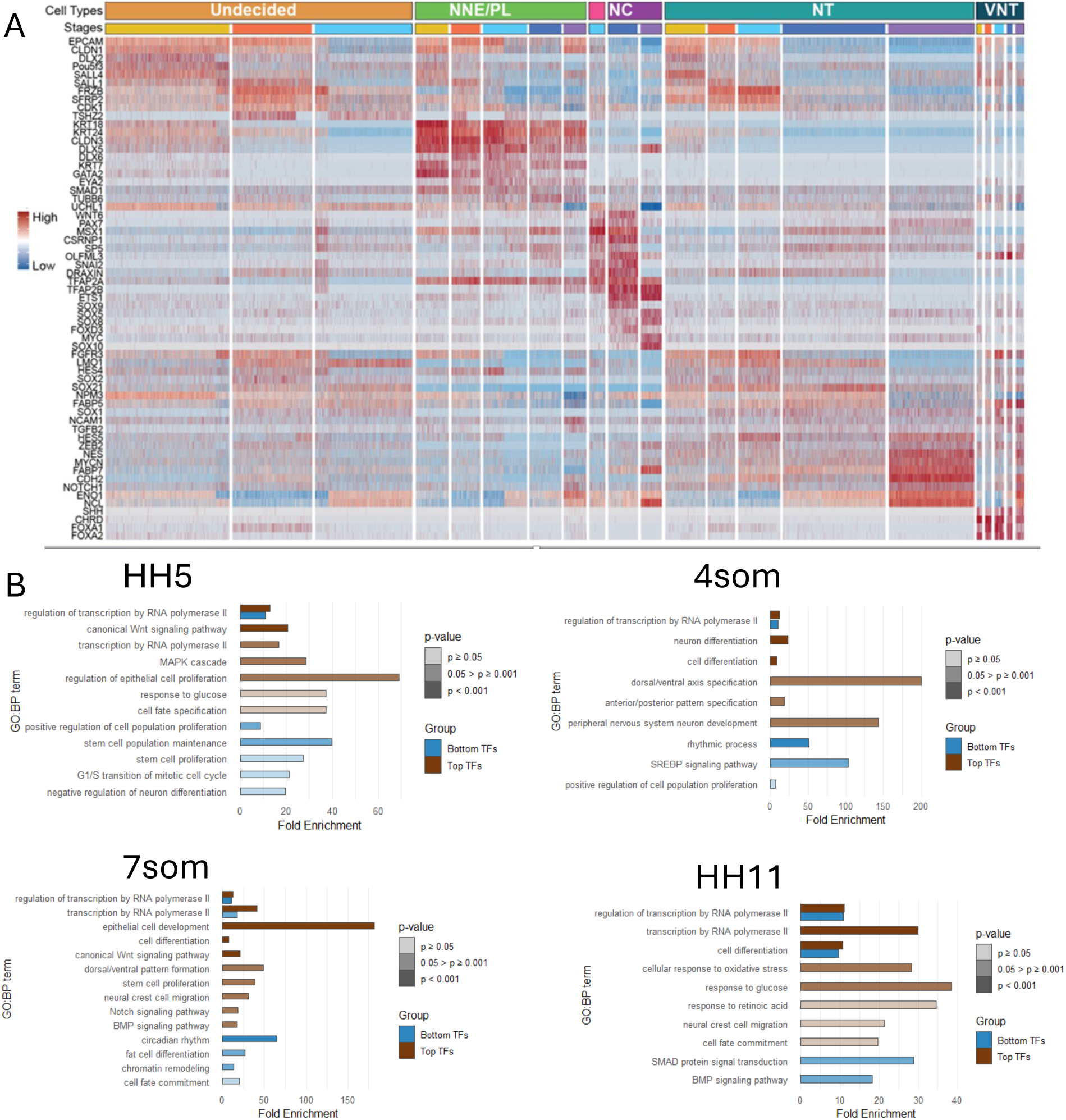
**A)** Heatmap showing top differentially expressed genes per 21 unique idents. Full list of genes is in supplemental Table 2 (minimum percent = 0.25, log fold-change threshold = 0.2). Top gene ontology terms consisting of genes with top and bottom (n= 30 each) gene activity scores at **B)** HH5, **C)** 4somite **D)** 7 somite **E)** HH11. Bars show fold enrichment, with significance represented as –log10(Benjamini FDR).

**Supplemental Fig 3.**
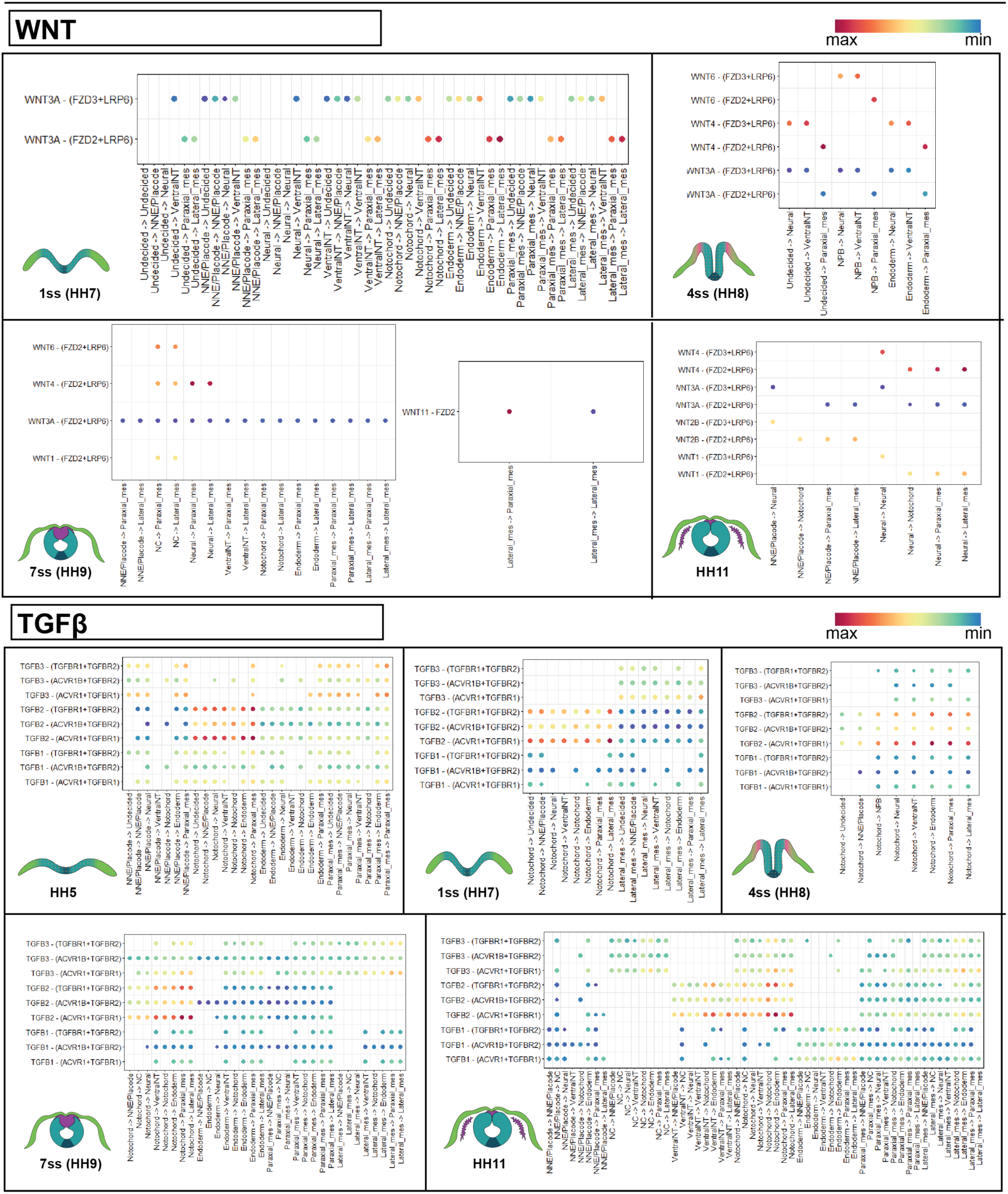
Individual Cell Chat ligand-receptor pair interaction predictions for Wnt (1som-HH11) and TgfB-signaling (HH5-HH11).

**Supplemental Fig 4.**
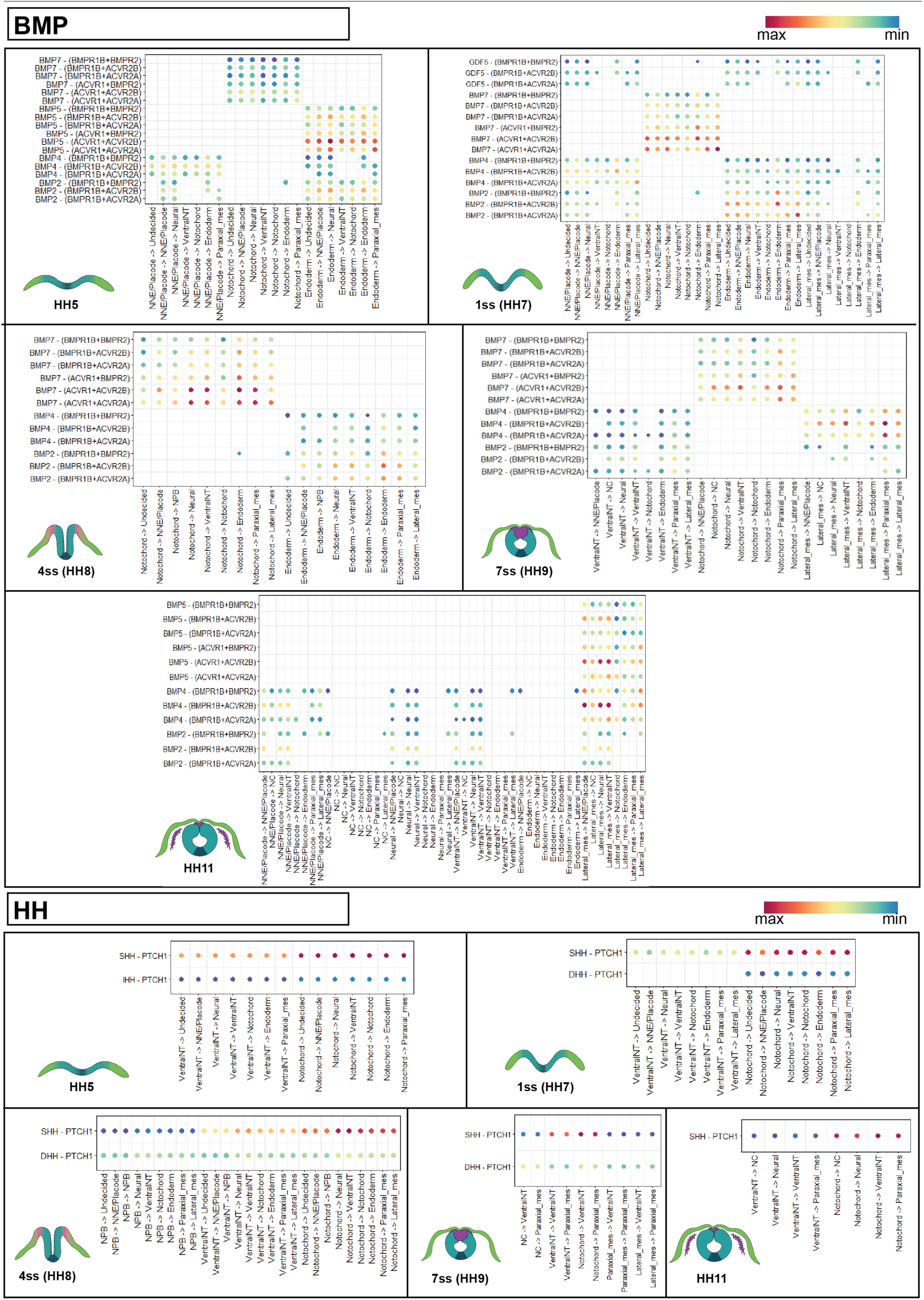
Individual Cell Chat ligand-receptor pair interaction predictions for BMP and Hedgehog-signaling pathways (HH5-HH11).

**Supplemental Fig 5.**
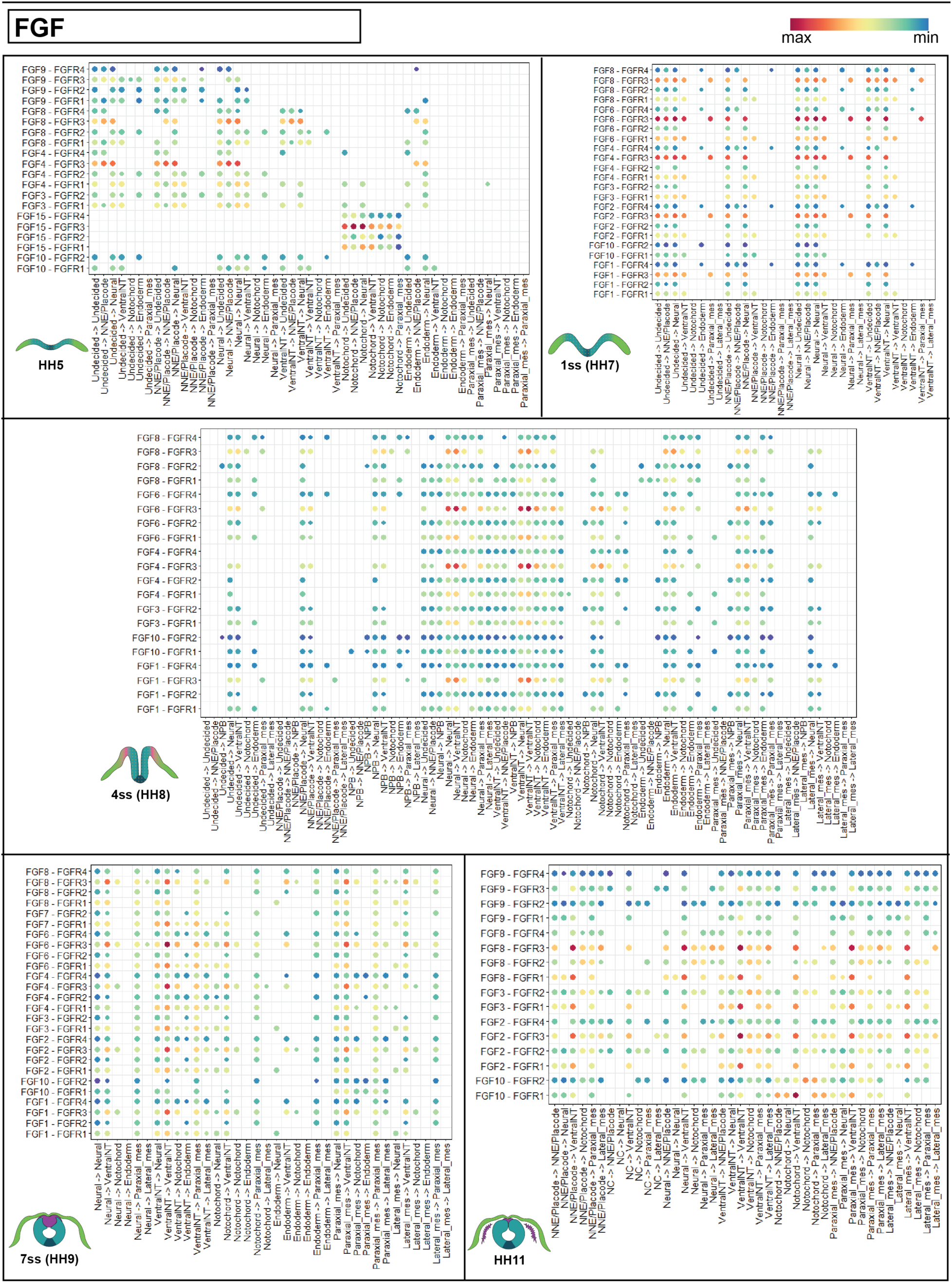
Individual Cell Chat ligand-receptor pair interaction predictions for the FGF signaling pathway (HH5-HH11).

**Supplemental Figure 6.**
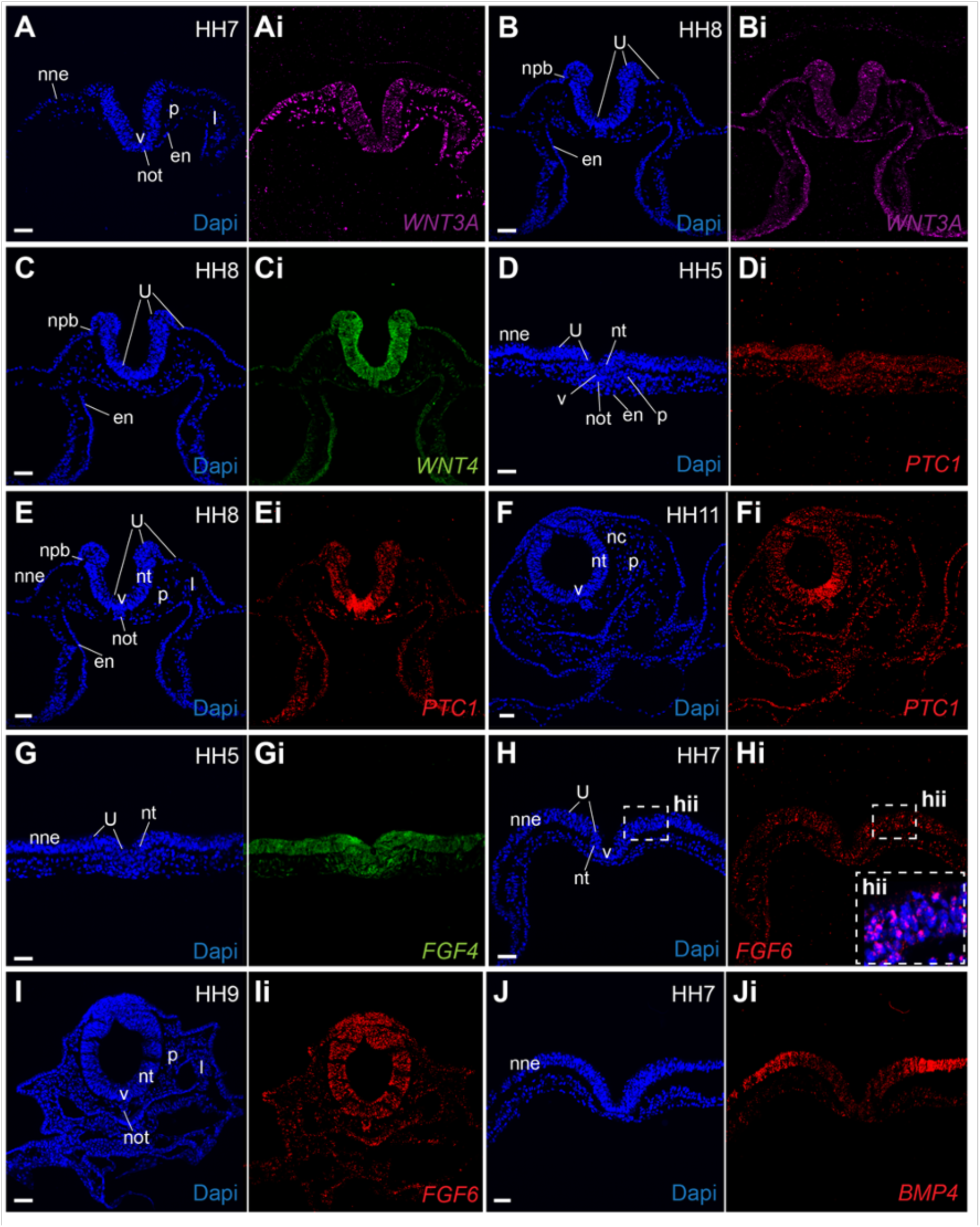
HCR-amplified fluorescent *in situ* hybridization-based validation of Cell Chat predictions not found in the literature. **A)** *Wnt3A* at HH7 **B)** *Wnt3A* at HH8 **C)** *Wnt4* at HH8, **D)** *Ptc1* at HH5, **E)** *Ptc1* at HH8, **F)** *Ptc1* at HH11 **G)** *Fgf4* at HH5, **H)** *Fgf6* at HH7 **I)** *Fgf6* at HH9 **J)** *Bmp4* at HH7. Scale Bar 50µm.

**Supplemental Fig 7.**
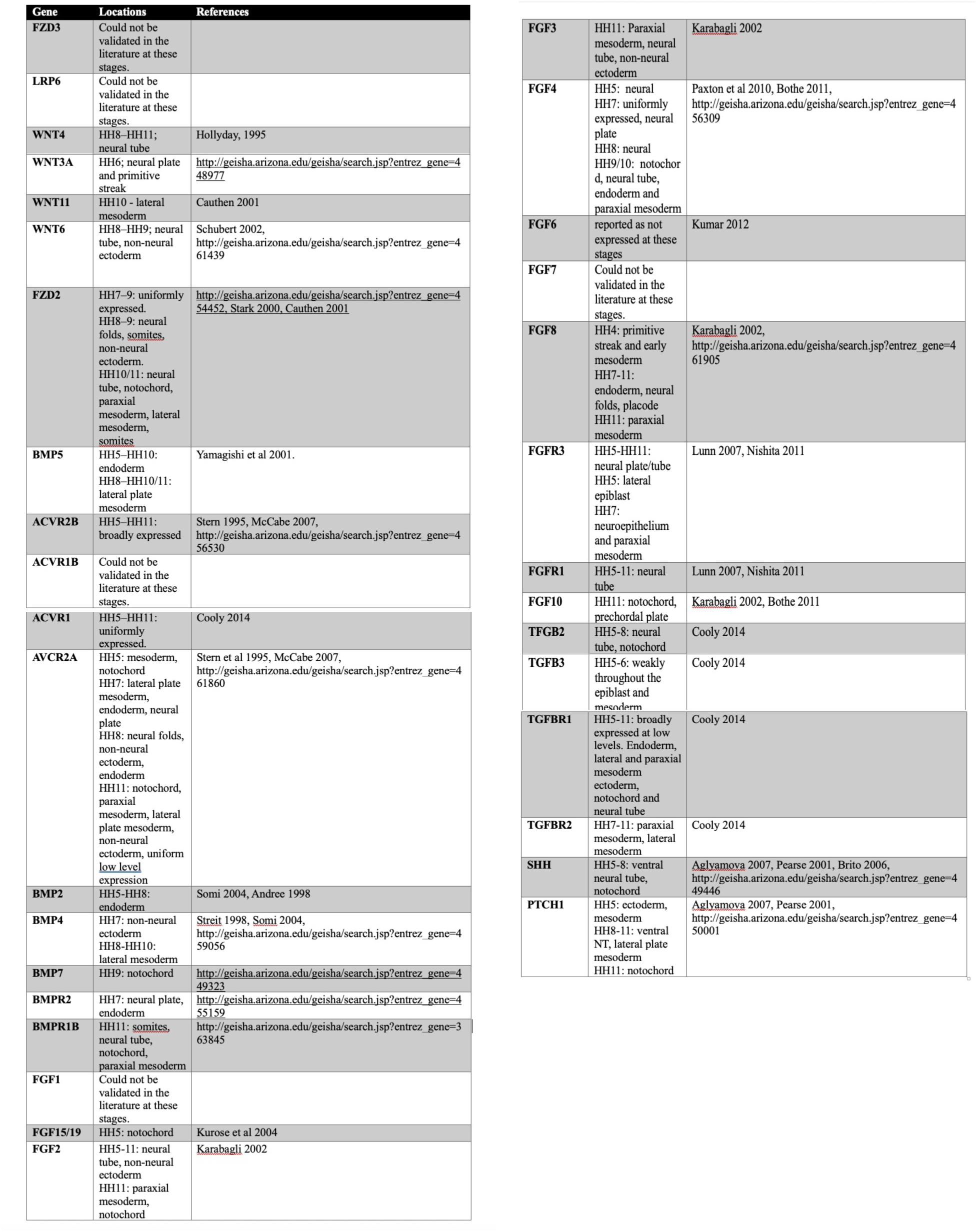
Literature sources for published *in situ* hybridization data that validates Cell Chat predictions.

